# A structure-derived mechanism reveals how capping protein promotes nucleation in branched actin networks

**DOI:** 10.1101/2021.03.15.435411

**Authors:** Johanna Funk, Felipe Merino, Matthias Schaks, Klemens Rottner, Stefan Raunser, Peter Bieling

## Abstract

Heterodimeric capping protein (CP/CapZ) is an essential factor for the assembly of branched actin networks, which push against cellular membranes to drive a large variety of cellular processes. Aside from terminating filament growth, CP stimulates the nucleation of actin filaments by the Arp2/3 complex in branched actin networks through an unclear mechanism. Here, we report the structure of capped actin filament barbed ends, which reveals how CP not only prevents filament elongation, but also controls access to both terminal filament subunits. In addition to its primary binding site that blocks the penultimate subunit, we find that the CP sterically occludes the central interaction site of the terminal actin protomer through one of its C-terminal “tentacle” extensions. Deletion of this β tentacle only modestly impairs capping. However in the context of a growing branched actin network, its removal potently inhibits nucleation promoting factors (NPFs) by tethering them to capped filament ends. End tethering of NPFs prevents their loading with actin monomers required for activation of the Arp2/3 complex and thus strongly inhibits branched network assembly both in cells and reconstituted motility assays. Our results mechanistically explain how CP couples two opposed processes –capping and nucleation– in branched actin network assembly.

## Introduction

Heterodimeric capping protein (CP/CapZ) is an evolutionarily ancient actin regulator found in nearly all eukaryotic organisms and cell types (Cooper and Sept, 2008; Edwards et al., 2014; Prostak et al., 2020; Rivero and Cvrcková, 2007) that controls the fate and interactions of actin filament ends in a variety of cellular contexts. Comprised of two closely related α and β subunits, it forms a constitutive heterodimer that tightly binds to the barbed end of actin filaments to terminate their growth. Within sarcomeres, it caps the barbed ends of filaments in the z-disk. In non-muscle cells, CP collaborates with a core set of proteins –profilin, the Arp2/3 complex and a membrane-localized nucleation promoting factor (NPF) of the WASP protein family– to create dense, polarized networks of short and branched filaments (Akin and Mullins, 2008; Cameron et al., 2001; Loisel et al., 1999). These branched actin networks generate pushing forces to move cellular membranes (Bieling et al., 2016; Mueller et al., 2017; Papalazarou and Machesky, 2020) in a large number of cellular processes ranging from cell motility (Wu et al., 2012), endocytosis (Senju and Lappalainen, 2019), phagocytosis (Jaumouillé and Waterman, 2020), autophagy (Kast and Dominguez, 2017) to cell-cell adhesion (Efimova and Svitkina, 2018). CPs central role in branched network assembly is commonly thought to simply result from its eponymous biochemical activity – to cap actin filament barbed ends and inhibit the addition of further subunits (Blanchoin et al., 2014; Edwards et al., 2014; Isenberg et al., 1980; Schafer et al., 1996). However, in vitro reconstitutions of branched network assembly suggest a more complex function (Akin and Mullins, 2008).

Capping protein is known to contribute to branched actin network assembly in several ways. First, it restricts the growth of filaments to short length, which prevents filament buckling for efficient force generation (Cameron et al., 2001; Vinzenz et al., 2012; Ydenberg et al., 2011). Second, it rapidly stops the non-productive elongation of filaments distant from the cell membrane. This generates a requirement for continuous nucleation of new filament branches via the Arp2/3 complex, which is in turn activated by a membrane-bound NPF. Confinement of Arp2/3 activation to the membrane is therefore responsible for polarized network growth (Achard et al., 2010; Akin and Mullins, 2008). In addition to these established functions directly related to the termination of filament growth, CP has been shown to stimulate nucleation of new filaments by the Arp2/3 complex. Perturbing CP activity either in vivo or in reconstituted actin motility results in similar effects that are consistent with it acting as a positive regulator of Arp2/3-dependent branching nucleation (Akin and Mullins, 2008; Hug et al., 1995; Loisel et al., 1999; Mejillano et al., 2004; Mullins et al., 2018). However, the responsible biochemical mechanism has remained enigmatic.

Branching nucleation itself is a multi-step process that requires the transient formation of a higher-order complex between the Arp2/3 bound to the side of an actin mother filament and a NPF that is loaded with an actin monomer bound to its WH2 domain (Rotty et al., 2013). It is presently unclear, which steps in this process determine the rate of filament nucleation within growing actin networks. The activity of CP seems to be a key factor (Akin and Mullins, 2008), but we currently do not understand how CP might either directly or indirectly influence the Arp2/3 complex, its upstream NPF or any intermediates formed between the two during nucleation. Prior structural and biochemical work on CP provides little insight into this crucial question (Edwards et al., 2014). One potential hint comes from the observation that each CP subunit possesses a C-terminal helical extension known as tentacles, which share sequence homology with the actin-binding NPF WH2 domain. Based on a low-resolution structure of capped skeletal muscle actin filaments (Narita et al., 2006) it has been proposed that most of the interaction occurs between the body of CP, which, together with its α tentacle, contacts the penultimate actin protomer. In addition, the β tentacle constitutes a potential secondary actin-binding region that has been speculated to bind to the last actin subunit (Edwards et al., 2014; Kim et al., 2010; Narita et al., 2006; Wear et al., 2003). While this is indeed the case in the homologous complex found within the dynactin complex in which CP caps an short Arp1 filament (Urnavicius et al., 2015), the structural details of capped actin filaments are still unknown. As a consequence, we understand very little how CP might influence Arp2/3 activity.

One conceptual model of why CP stimulates nucleation in branched networks is offered by the so-called monomer gating hypothesis (Akin and Mullins, 2008; Mullins et al., 2018). Key to monomer gating is a presumed kinetic competition between growing filament ends and the Arp2/3 complex for a limiting flux of actin monomers already in complex with the membrane-bound NPFs. As polymerizing filament ends become more numerous through branching, they should consume larger amounts of NPF-bound monomers, which might reduce their partitioning into nucleation via the Arp2/3 complex. While this model offers a conceptual explanation for the nucleation promoting function of CP, it lacks direct experimental evidence. Indeed, alternative mechanisms based on the local depletion of *soluble* actin monomers have also been proposed (Wang and Carlsson, 2015).

We are currently lacking a comprehensive mechanism that explains and unifies the distinct roles of CP based on its interactions with the other core components of the branched actin network. To obtain such an understanding, we used electron cryo-microscopy (Cryo-EM) to determine the structure of the mammalian CP heterodimer bound to non-muscle actin filaments. We find that in addition to its main binding site located in between the two last actin subunits, CP contacts the barbed end groove of the ultimate actin subunit via the tentacle extension of its β subunit similar to -and mutually exclusive with binding of NPF WH2 domains. Using single molecule TIRFM assays and reconstitution of branched network assembly, we demonstrate that this type of “masking activity” of the CP β tentacle is dispensable for barbed end capping, but required for sustained nucleation of new filaments in branched actin networks by NPF-activated Arp2/3 complexes *in vitro*. In line with this, we find that the CP β tentacle is selectively required for efficient lamellipodial protrusion of mammalian cells and the localization of CP to the leading edge. In the absence of the CP β tentacle, NPFs tether to capped actin filament ends in a nucleation-deficient state. Our results reveal an unanticipated mechanism at the core of the branched actin network engine that couples the creation of polymerizing filaments by Arp2/3 to their elimination through capping protein.

## Results

### The structure of CP-bound actin filament barbed ends

The ends of actin filaments have emerged as important regulatory sites, to which a diverse set of regulators bind to control filament dynamics. Despite their obvious importance, there are -with a single exception (Shaaban et al., 2020)- no high-resolution structures of filament ends available. Because filament elongation is kinetically far more favorable than nucleation, the exceedingly sparse density of ends in common actin filament preparations has prevented single particle analysis until now. To overcome this fundamental limitation, we established a general procedure to generate short actin filaments amenable to single-particle Cryo-EM analysis (Figure 1A). To this end, we premixed CP with actin in buffers of low ionic strength containing Ca^2+^, which keep it from polymerizing. We then triggered rapid filament nucleation by salt addition. Under these conditions, and in the absence of monomer-binding proteins such as profilin, CP potently nucleates filaments that slowly elongate from the pointed end (Cooper and Pollard, 1985). We stabilized these capped filament stubs using phalloidin shortly after the initiation of polymerization. To separate stabilized filaments from residual soluble capping protein and actin, we subjected these reactions to size exclusion chromatography (Figure 1B). SDS PAGE revealed the presence of both CP and actin at roughly 1:20 stoichiometry (Supplemental Figure 1A) in the high molecular weight fractions (Figure 1B).

**Figure 1:**
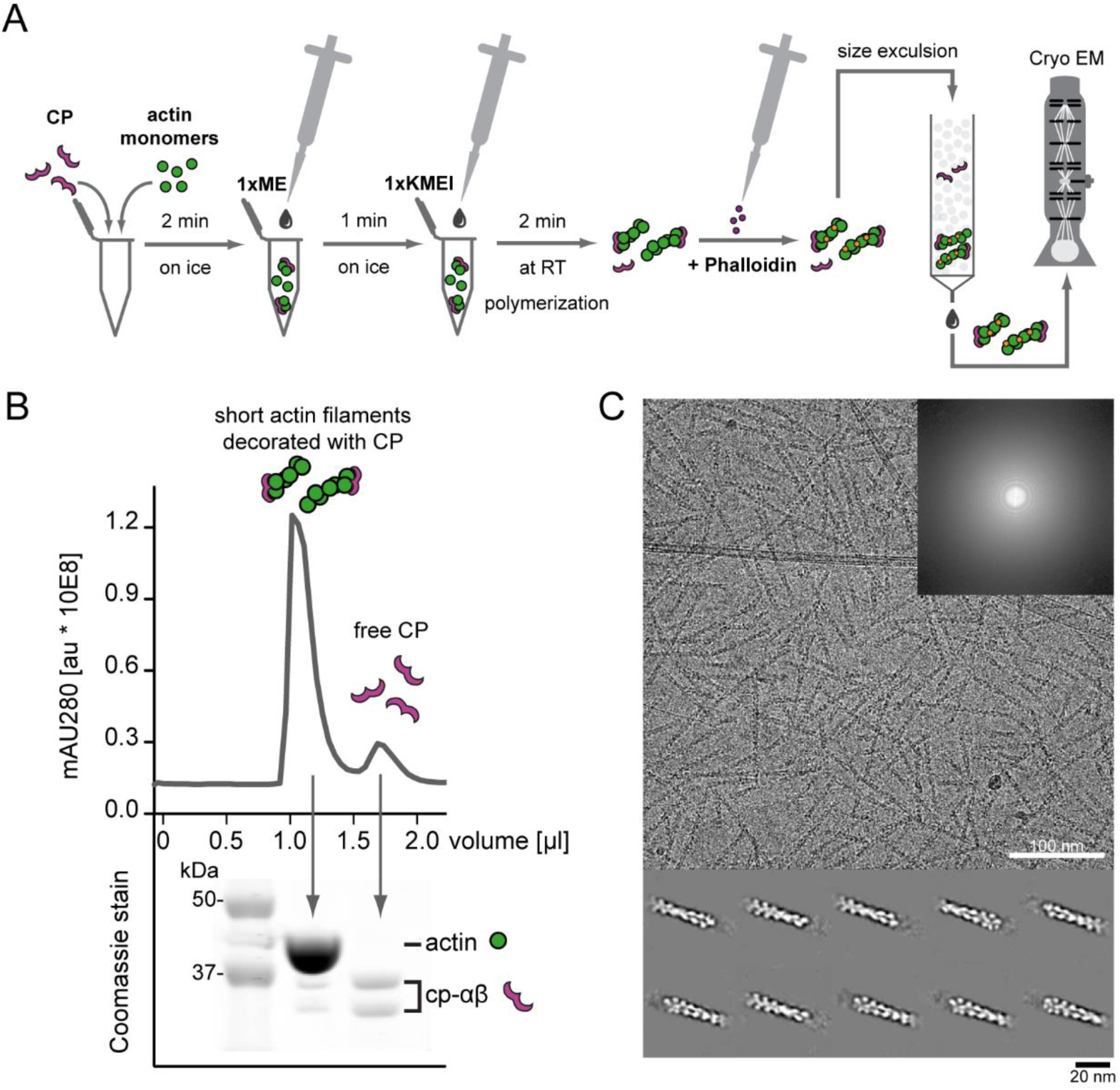
Preparation and visualization of CP-bound barbed ends by cryo-EM. (A) Workflow of capped filament preparation for cryo-EM. Monomers were mixed with CP followed by Ca^2+^-to-Mg^2+^ exchange and salt addition to trigger polymerization. Polymerization was arrested by phalloidin addition and capped filaments were separated from free CP by size-exclusion chromatography (SEC) and then visualized by cryo-EM. (B) Isolation of short, capped actin-filaments from free CP by SEC. Top: chromatography profile. Bottom: Corresponding Coomassie-stain with bands representing actin or α and β-capping protein as indicated (Supplemental Figure 1). (C) Representative micrograph of vitrified capped filaments on graphene oxide grids imaged at 1.5 µm defocus. The inset shows the corresponding power spectrum for the image. Bottom: Class averages showing densities for capping protein bound to respective filament barbed ends

Cryo-EM images of samples from these fractions revealed the presence of densely packed filaments of near-uniform lengths of 80-120 nm; Figure 1C) Using these samples and an iterative classification strategy we determined a Cryo-EM structure of CP-bound cytoplasmic actin barbed ends to an overall resolution of 3.8 Å (Figure 2A, Supplemental Figure 2A-B, Table S2). Local resolution ranges from 3-5 – 7.5 Å, with most actin monomers resolved better than 4 Å and CP to 6 - 7 Å (Supplemental Figure 2B).

**Figure 2:**
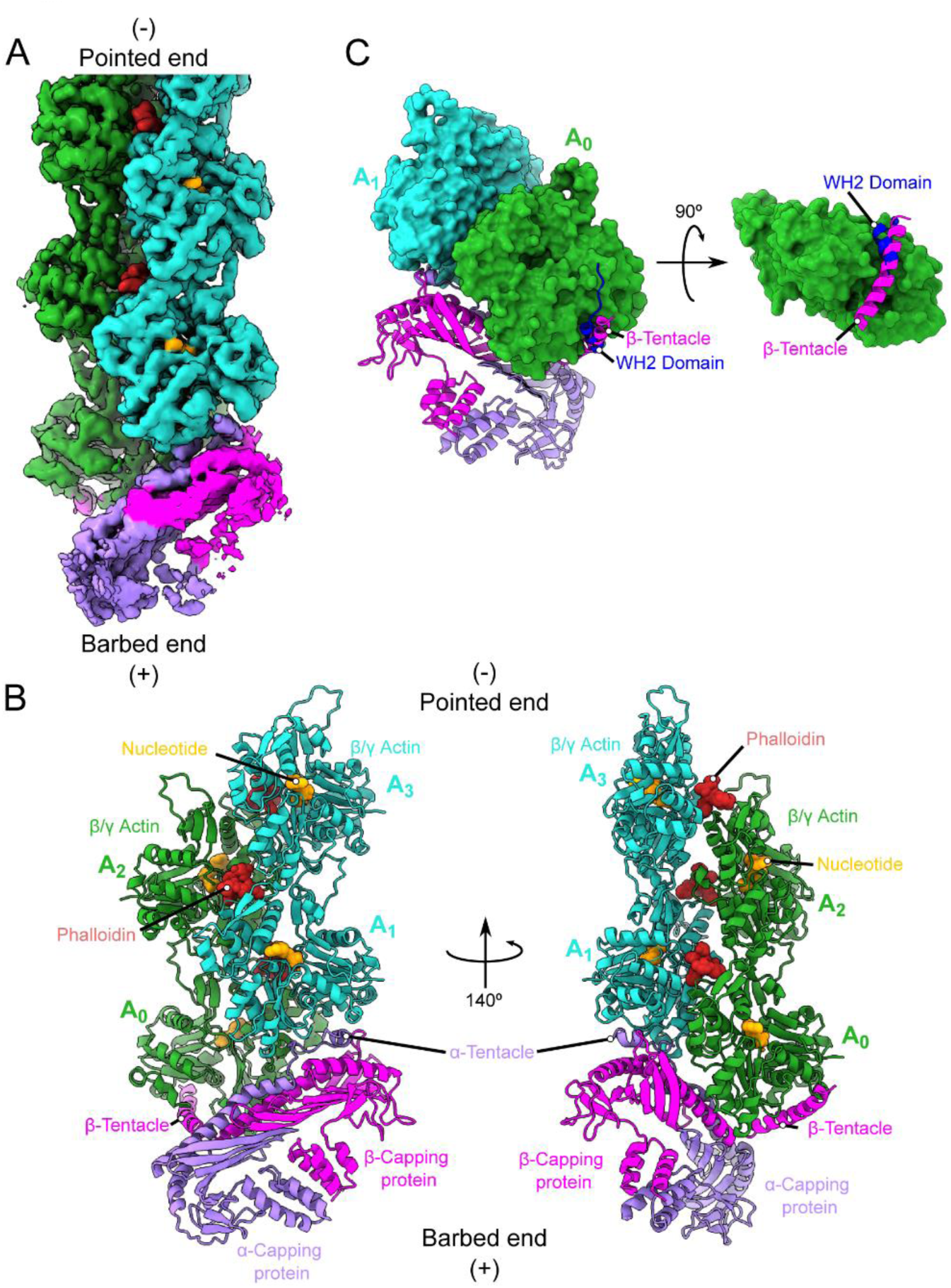
Cryo-EM structure of CP-bound barbed ends. (A) Overview of the Cryo-EM map of capped F-actin. The density has been sharpened and denoised using deepEMhancer (Sanchez-Garcia et al., 2020) (B) Atomic model of CP bound to the barbed end. While the penultimate protomer interacts extensively with CP, the terminal protomer is bound mostly by the β tentacle. We highlight the position of the bound small molecules. Actin subunits are labeled A_0_ to A_3_ starting from the terminal actin protomer. (C) The structure of WASP-bound G-actin (Chereau et al., 2005, PDBID 2A3Z) was superimposed onto the terminal protomer of the complex. The β tentacle occupies the binding pocket that a WH2 domain would need to bind the terminal protomer. Actin monomers are shown as surface, while the tentacle and the WH2 helices are shown as cartoons.

As expected, the structure of cytoplasmic actin filament closely resembles the previously determined structures of skeletal actin (Merino et al., 2018; Oda et al., 2009). We observed clear density for phalloidin in all the expected binding sites (Pospich et al., 2020), with the exception of the cleft between the ultimate and penultimate actin subunits (Supplemental Figure 2C). This is likely due to the absence of the missing third actin protomer required to form the full binding site. Consistent with our previous work (Merino et al., 2018), we find the D-loop in a closed state. Due to the comparably high resolution within the nucleotide-binding site of actin, we could directly determine the identity of the nucleotide bound to each subunit. Consistent with the early addition of phalloidin during the polymerization reaction, most subunits show clear density for inorganic phosphate (P_i_) (Supplemental Figure 2D). Interestingly, the two terminal subunits lack this density, suggesting that they are in an ADP-bound state. Whether this was due to the direct interaction with CP favoring Pi release or simply the result of our filament preparation procedure, in which these two subunits are the “oldest” in the filament as they form with CP the nucleus for polymerization is unclear.

As expected, the primary interaction between CP and the filament occurs at the barbed end face of the penultimate actin subunit (Figure 2B). The core of CP together with the α tentacle block all of the exposed barbed end surface of the penultimate actin protomer, with the α tentacle deeply wedged in a hydrophobic cleft at the barbed end of this subunit. In contrast, CP makes few contacts with the ultimate protomer. Minor contacts are formed between CPs core and the inner side of the ultimate protomer. Notably, a clear density occupies the hydrophobic cleft at the barbed end of this subunit, which can be unequivocally assigned to the β tentacle (Figure 2B, C). The density in this region was slightly weaker, but comparable to the body of the terminal actin protomer (Figure 2A), suggesting that the β tentacle remains mostly docked to actin and but retains some flexibility. The overall interaction between CP and the actin barbed end is overall very similar to what has been observed in the dynactin complex (Urnavicius et al., 2015), with both tentacles occupying equivalent positions in contacting one end of the Arp1 filament. The placement of the β tentacle has only been speculated from a prior low-resolution structure of capped skeletal actin (Narita et al., 2006) and agrees with prior computational models and mutational analysis (Kim et al., 2010; Wear et al., 2003). As predicted previously (Kim et al., 2010), this secondary barbed end binding site conspicuously overlaps with and mirrors the interaction between WH2 domain-containing proteins of the WASP family and actin monomers (Figure 2C). The strong conservation of the β tentacle among even evolutionarily distant CP family members suggests that this specific type of interaction serves an important function.

### Deletion of the CP β tentacle modestly affects barbed end capping kinetics

To understand the role of this conserved secondary actin-binding site in CP, we removed the last 23 amino acids of the beta subunit (CPαβΔ23) comprising the full β tentacle (Figure 3A) and studied the effect on filament barbed end association and dissociation kinetics by TIRFM assays in vitro (Figure 3). First, we measured the rate of CP association to growing, surface-tethered actin filaments at the single molecule level (Figure 3B, C). Observed capping rates (Supplemental Figure 3) were plotted against the total CP concentration and linear fits to the data yielded association rate constants (k_on_, Figure 3D). In a second line of experiments, we probed CP dissociation from the barbed ends of surface-tethered filaments by washout assays (Figure 3E, F). Due to the extraordinary long lifetime of CP at the barbed end (>10 min), we could not visualize single CP molecules directly but inferred dissociation indirectly from the moment of filament regrowth (Figure 3E, F). Mono-exponential fits to the dwell time courses yielded dissociation rate constants (k_off_, Figure 3G). In overall agreement with prior bulk measurements (Kim et al., 2004; Wear et al., 2003), we found that the deletion of the entire CP β tentacle reduced the affinity for the barbed end only by about one order of magnitude. This was the result of an equally moderate (3-fold) change in association and dissociation rate constants. The moderate drop in affinity suggests that engagement of this secondary binding site is not essential for filament capping.

**Figure 3:**
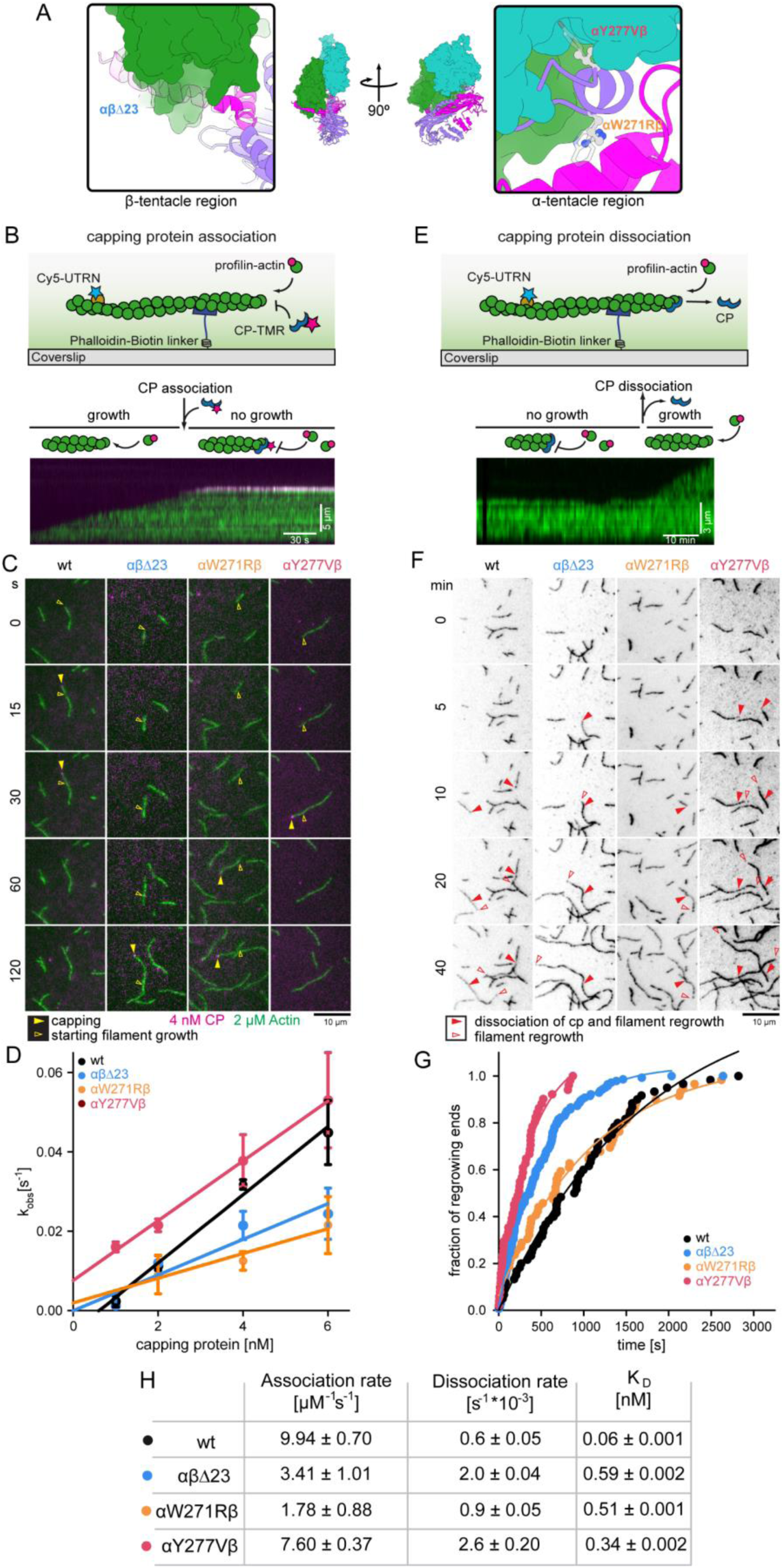
The CP β tentacle is dispensable for capping of single actin filaments. (A) Overview of the location of CP mutations. Colors are as in Fig 2. Insets highlight the region of the β tentacle deleted (green), and the color scale on the actin surface depicts increasing hydrophobicity from gray to white to yellow. (B) Scheme of single molecule TIRFM assays (top). Individual surface-attached filaments visualized by Cy5-UTRN261N (10nM, green) grow from profilin-actin (2 μM) in the presence of TMR-CP (4 nM, magenta). CP association results in the termination of filament growth as visualized in the kymograph (bottom). (C) TIRFM time lapse images of filament elongation (green) in presence of wt or mutant capping protein (magenta) as indicated under conditions as in (B). Empty yellow arrows indicate the initial position of the barbed end and filled yellow arrows the position at the moment of CP association. (D) Observed reaction rates (k_obs_) for capping protein wt and mutants as a function of CP concentration (Supplementary Figure 3). Association rate constants (k_on_) are calculated from linear fits to the data. (E) Scheme of actin filament barbed end re-growth after dissociation of CP via TIRFM (top). Single capped filaments were visualized by Cy5-UTRN261N (10 nM, green) in the presence of profilin-actin (2 uM) and myotrophin/V1 (20 nM) after CP washout. CP dissociation leads to growth of the filament visualized in the kymograph (bottom). (F) TIRFM time lapse images of filaments following the washout of either wt CP or mutants, as indicated. Conditions are as in (E). Filled red arrows indicate the positions of barbed ends at the moment of CP dissociation in each case, and empty red arrows follow the positions of (re-) growing barbed ends. (G) Time traces of the fraction of uncapped barbed ends after washout of either wt CP or mutants, as indicated, at t=0. Dissociation rate constants (k_off_) were derived from mono-exponential fits (see Material and methods). (H) Summary table of association (k_on_), dissociation rate constants (k_off_) and equilibrium dissociation constants (K_D_, calculated from the former rate constants) for the interaction of CP (wt or mutants as indicated) with actin filament barbed ends. Errors are SD of the mean values from 3 independent experiments (k_off_) or the SEM of the linear fit (k_on_).

The dwell time of CP at filament ends in vitro exceeds its lifetime in cellular actin networks by two orders of magnitude (Iwasa and Mullins, 2007; Lai et al., 2008; Miyoshi et al., 2006). This is in part the result of its active removal from filaments ends via twinfilin (Hakala et al., 2021; Mwangangi et al., 2021). It is reasonable to assume therefore that its association rate should be functionally more relevant compared to its exceedingly slow in vitro dissociation rate. Even in the absence of its β tentacle, CP still dwells at filament ends far longer than it should be necessary to carry out its capping function at the leading edge of motile cells. To control for the moderate contribution of the β tentacle to barbed end capping kinetics and to uncouple this effect from a potential unrelated secondary function, we designed weakening mutants distant from β tentacle within the primary barbed end binding site of CP (Figure 3A, see Materials and Methods), We identified two single site mutants within the CP α subunit (αW271Rβ and αY277Vβ) that altered the thermodynamics and kinetics of barbed end binding in a fashion highly similar to β tentacle removal (Figure 3D,G,H). We therefore refer to these two variants as “kinetics-mimicking” mutants hat closely resemble the effect of the β tentacle deletion on the single filament capping rates.

### Deletion of the CP β tentacle drastically inhibits reconstituted branched actin network assembly

Because the deletion of the CP β tentacle results in only mild changes in the kinetics of capping, we wondered how this relates to its potentially more complex role in regulating branched actin network dynamics. To address this question, we assembled branched actin networks from polystyrene beads that were coated with a minimal Arp2/3-activating fragment of the NPF protein WAVE1 (WAVEΔN, (Bieling et al., 2018)). We initiated network assembly by the addition of Arp2/3 complex (100nM), CP (wildtype or mutants, 100nM), and profilin–actin (5 µM); conditions previously shown to promote rapid network growth (Akin and Mullins, 2008; Bieling et al., 2016). To visualize the kinetics of all biochemical processes contributing to the assembly of the actin network-actin polymerization, Arp2/3-driven nucleation and CP-mediated capping-we included a fluorescently labeled filament binding probe (Alexa488-Lifeact, (Burkel et al., 2007; Funk et al., 2019)) and used a fraction of Arp2/3 and CP labeled with matching fluorescent dyes (Figure 4A). We kinetically arrested network assembly very soon (4 min) after the initiation of growth by adding a 3-fold molar excess of Latrunculin B and phalloidin, which stabilizes existing actin filaments and prevents the nucleation of new ones (Akin and Mullins, 2008). Upon kinetic arrest, we immediately proceeded to multi-color imaging of the actin networks (Figure 4B).

**Figure 4:**
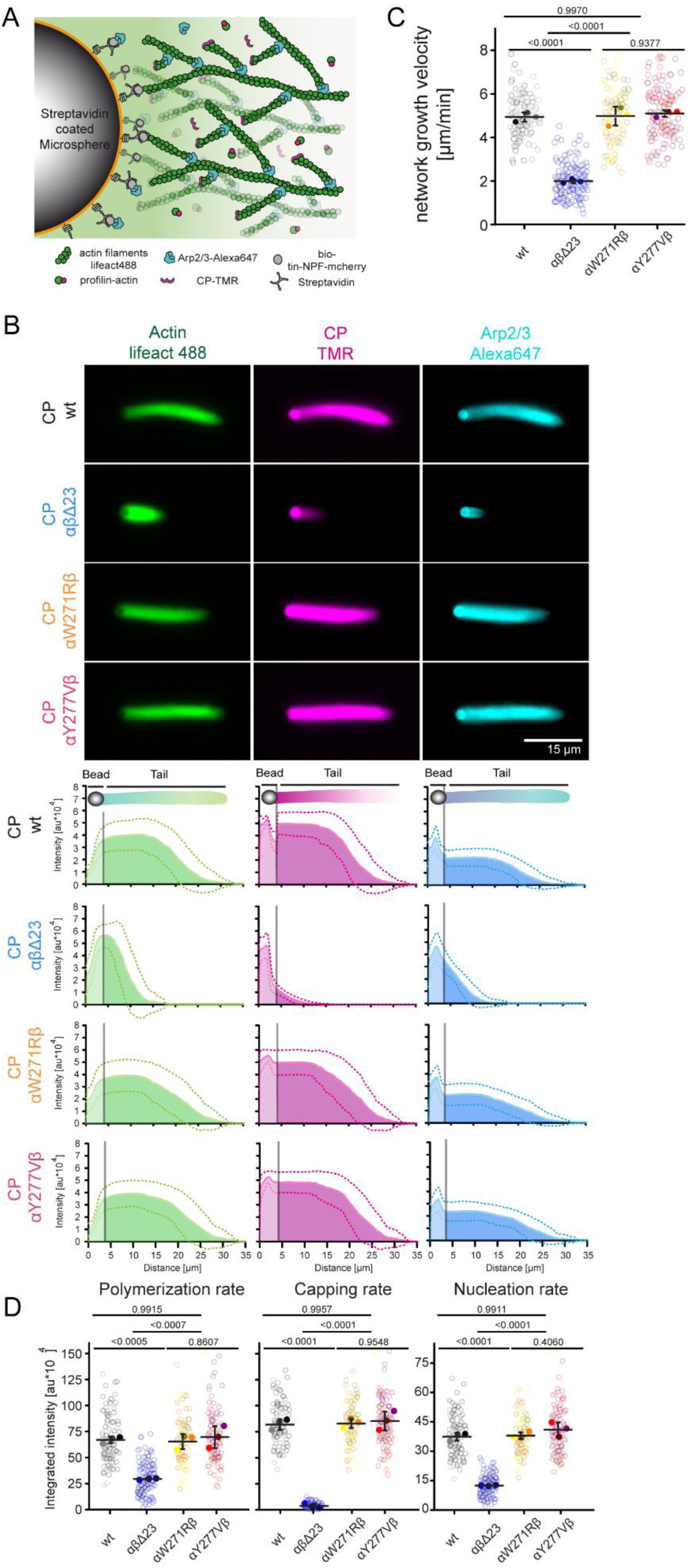
The CP β tentacle is essential for reconstituted branched actin network assembly. (A) Scheme of bead-motility-assays. Dendritic actin networks are assembled from polystyrene microspheres (Ø = 3 μm) coated with WAVE1ΔNvia biotin-streptavidin linkages. Dense branched actin networks assemble from these beads and incorporation of all constituting components can be visualized via matching fluorescent labels as indicated. (B) Top: Epifluorescence mages of dendritic actin networks grown from WAVE1ΔN-coated microspheres using 5 μM profilin–actin, 100 nM CP (either wt or mutants as indicated, 20 % TMR-labeled), 100 nM Arp2/3 (20 %-Alexa647 labeled). Reactions were kinetically arrested after 4 min using phalloidin (15 μM), Latrunculin-B (15 μM) and Alexa488-Lifeact (15 nM) to visualize filamentous actin. Bottom: Quantification of the averaged intensity profiles for the indicated dendritic network components. (C) Plots of network growth velocities of dendritic actin networks grown in presence of either wt or mutant CP. (D) Plots of the polymerization rates, the capping rates and the nucleation rates of dendritic actin networks grown in presence of either wt or mutant CP (see Material and Methods). Quantifications were done for N ≥ 25 tails from 3 independent experiments, error indicator = SD of the mean. P-values (see C and D) were derived from ANOVA Tukey tests.

We observed that reactions with CP lacking the β tentacle generated actin networks with drastically altered assembly dynamics compared to reactions with wildtype CP (Figure 4B, top). Averaged intensity profiles showed that networks grown in the presence of CPαβΔ23 contained less polymerized actin and drastically reduced Arp2/3 and CP amounts (Figure 4B, bottom). Relative to the wildtype CP control, these networks grew with about 3-fold reduced average velocities (Figure 4C). To directly quantify the rates at which actin, CP, and Arp2/3 join the growing network at the NPF-coated surface, we integrated the fluorescence intensity of each component in individual actin networks and divided these values by the reaction time (Figure 4D). This analysis revealed that the deletion of the CP β tentacle dramatically (20-fold, Supplemental Figure 4) reduced network incorporation rate of CP, much more than predicted by its modest defect in capping at the single molecule level (Figure 3H). Conversely, we also observed a drastic reduction in the rate of Arp2/3 network incorporation (Figure 4D). To test whether this reduced filament nucleation rate at the network level was simply a consequence of its modestly reduced capping rate, we turned to the CP mutants kinetically mimicking the β tentacle deletion (αW271Rβ and αY277Vβ). Decisively, we observed that both mutants were capable of generating branched actin networks with assembly kinetics nearly indistinguishable from wildtype CP (Figure 3B-D). We therefore concluded that the strongly diminished rates of network incorporation of both CP and Arp2/3 resulting from the CP β tentacle deletion were not due to its modestly altered capping rates that we observed at the single filament level. Instead, this implies that the CP β tentacle serves an important secondary function in branched network assembly not directly related to the termination of actin filament growth.

### Deletion of the CP β tentacle inhibits Arp2/3-dependent branching by tethering NPF proteins to capped filament ends through their WH2 domain

The CP β tentacle interacts with the central binding cleft of the terminal actin protomer at the barbed end in a manner that sterically prevents binding of other WH2 containing proteins such as NPFs (see above). We therefore hypothesized that capping protein controls the rate of Arp2/3-mediated branching at the level of the NPF as part of a feedback mechanism (Figure 5A): CP occupies both WH2 binding sites at the penultimate and terminal actin subunit to effectively mask the capped filament end from NPF binding. Loss of the β tentacle liberates one of these binding sites, which might drive the sequestration of proximal NPF WH2 domains. This type of tethering should therefore reduce the amount of NPF molecules loaded with actin monomers ready to activate the Arp2/3 complex.

**Figure 5:**
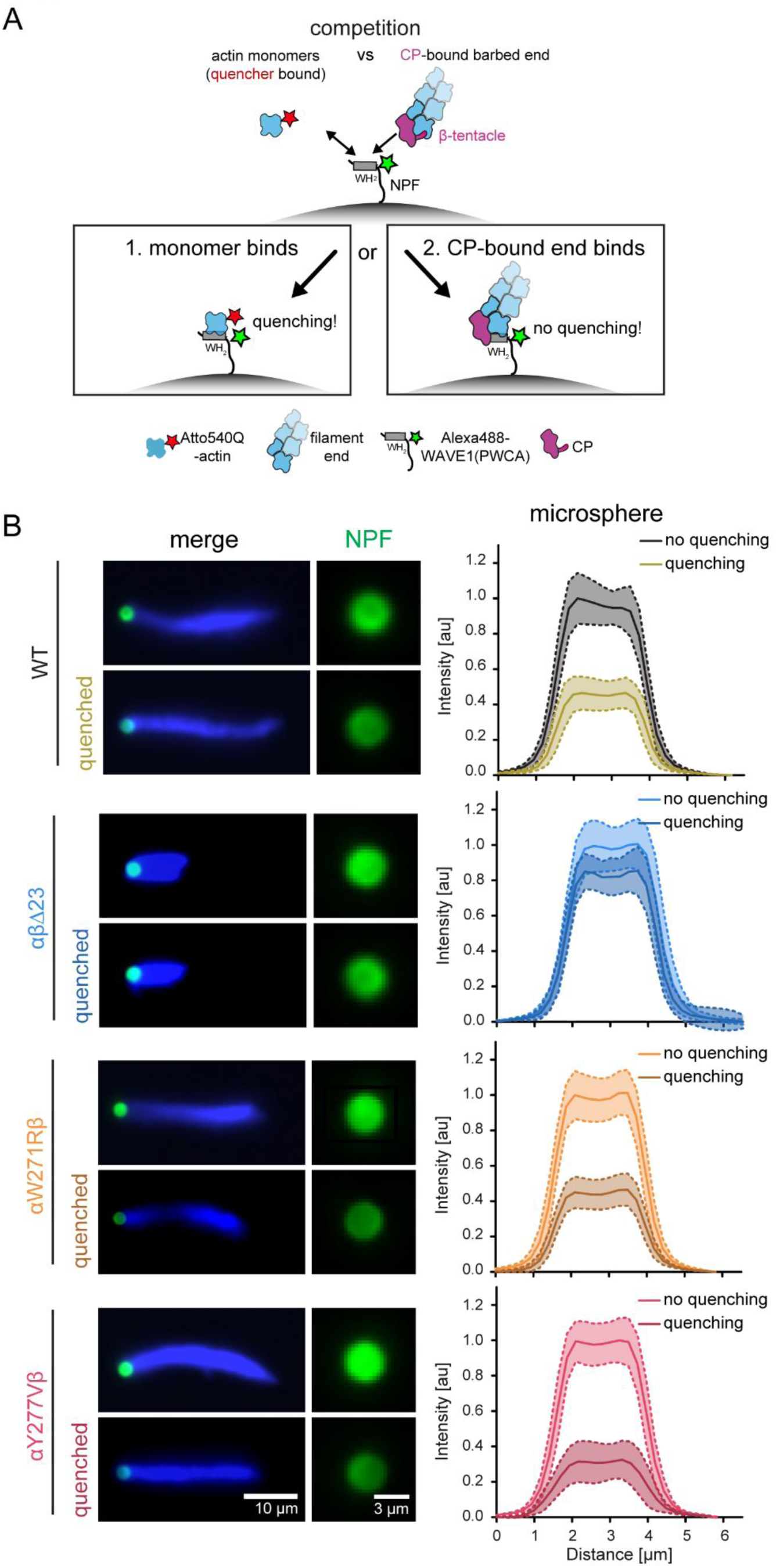
Loss of the CP β tentacle tethers NPFs to capped filament ends via their WH2 actin binding site. (A) Scheme of the FRET setup. Surface-bound WAVEΔN molecules are donor- (Alexa488-) labeled upstream of their WH2 site. NPF WH2 can either interact with quencher- (Atto540Q-) labeled actin monomers resulting in a decrease of donor fluorescence or unlabeled terminal protomers of filament barbed ends, resulting in no change in fluorescence. Terminal protomers are unlabeled, since quencher-labeled monomers are introduced only upon network arrest. Loss of the β tentacle vacates the WH2 binding site at the terminal protomer of capped ends, tipping the balance in favor of end-binding. (B) Left: Representative epifluorescence images of Alexa488-WAVEΔN-coated microspheres (green) and dendritic actin networks (visualized by Cy5-UTRN_261_, blue) 3 min after arrest before (top) and after (bottom) quenching with Atto540Q-labeled actin monomers. Protein concentrations were as in Figure 4 in the presence of wt or mutant capping proteins, as indicated. Right: Average intensities (error = SD) of the NPF-fluorescent signal intensity on the microsphere surface in the presence or absence of Atto540Q-labeled actin monomers (N ≥ 25 per condition).

To test this hypothesis, we employed a recently developed Förster resonance energy transfer (FRET) method to directly measure partitioning of WH2 domains between actin monomers and filament ends (Figure 3A, (Bieling et al., 2018)). To this end, we conjugated a fluorescent donor (Alexa488) to a site immediately upstream of the WH2 domain of WAVE1 and conjugated a non-fluorescent quencher (Atto540Q) to monomeric actin in a manner that does not affect their binding affinities (Bieling et al 2018, see Material and Methods). We then immobilized this labelled WAVE1 fragment on polystyrene beads and assembled branched networks in the presence of either wildtype CP or mutants. We added quencher-labeled and Latrunculin B-stabilized actin monomers at the moment of kinetic network arrest through Latrunculin B, phalloidin and myotrophin/V-1. The latter inhibitor preserves free barbed ends (Bieling et al., 2016). To minimize the impact of CP dissociation from capped ends after network arrest, we immediately proceeded to imaging and did not analyze NPF coated beads longer than 5 minutes after inhibitor and quencher addition. Alexa488-WAVEΔN-coated beads in physical contact with actin networks assembled from either wildtype or kinetics-mimicking (αW271Rβ and αY277Vβ) CP mutants displayed similarly strong (by 59-69%) quenching of NPF donor fluorescence, indicating that a majority of NPF molecules were able to bind quencher-labeled actin monomers from solution (Figure 5A,B). Strikingly, quenching was substantially diminished (to 14%) for Alexa488-WAVEΔN beads tethered to actin networks grown in the presence of CP lacking its β tentacle (Figure 5B). These results verify that the loss of the β tentacle locks the majority of NPF molecules in an inactive state tethered to capped filament ends. In summary, this strongly suggests that the conservation of this secondary end binding site in CP arises from the necessity to completely mask the capped barbed end from WH2 binding.

### The β tentacle is essential for CP localization to lamellipodial actin networks and efficient leading edge protrusion

To determine how our *in vitro* results translate to the function of CP in regulating branched actin network dynamics in mammalian cells, we generated engineered murine B16-F1 cell lines, in which expression of either one of the two CP subunits was genetically disrupted by CRISPR/Cas9-mediated genome editing. We first stably knocked out the single gene (*CapZb*) from which all three known CPβ isoforms are derived (Schafer et al., 1994). This successfully resulted in a clonal cell line devoid of detectable CPβ expression (CapZβ KO#10, Figure 6A). In parallel, we attempted to inactivate both genes (*CapZa1 and −2*) responsible for the expression of the two ubiquitous CPα isoforms (Cooper and Sept, 2008; Hart et al., 1997). The murine genome contains a third CPα gene (CapZa3), however the resulting protein is more divergent and only expressed in the male germ cell lineage (Cooper and Sept, 2008; Hurst et al., 1998). Simultaneous, CRISPR/Cas9-mediated targeting of *CapZa1* and *-2* did not yield clones entirely devoid of CPα, indicating that the CPα subunit might be essential for cell viability. Nonetheless, we were able to select a clone resulting from synchronous targeting of *CapZa1* and *−2* that showed markedly reduced total CPα expression levels, with the CPα1 isoform being virtually absent (Figure 6A).

**Figure 6:**
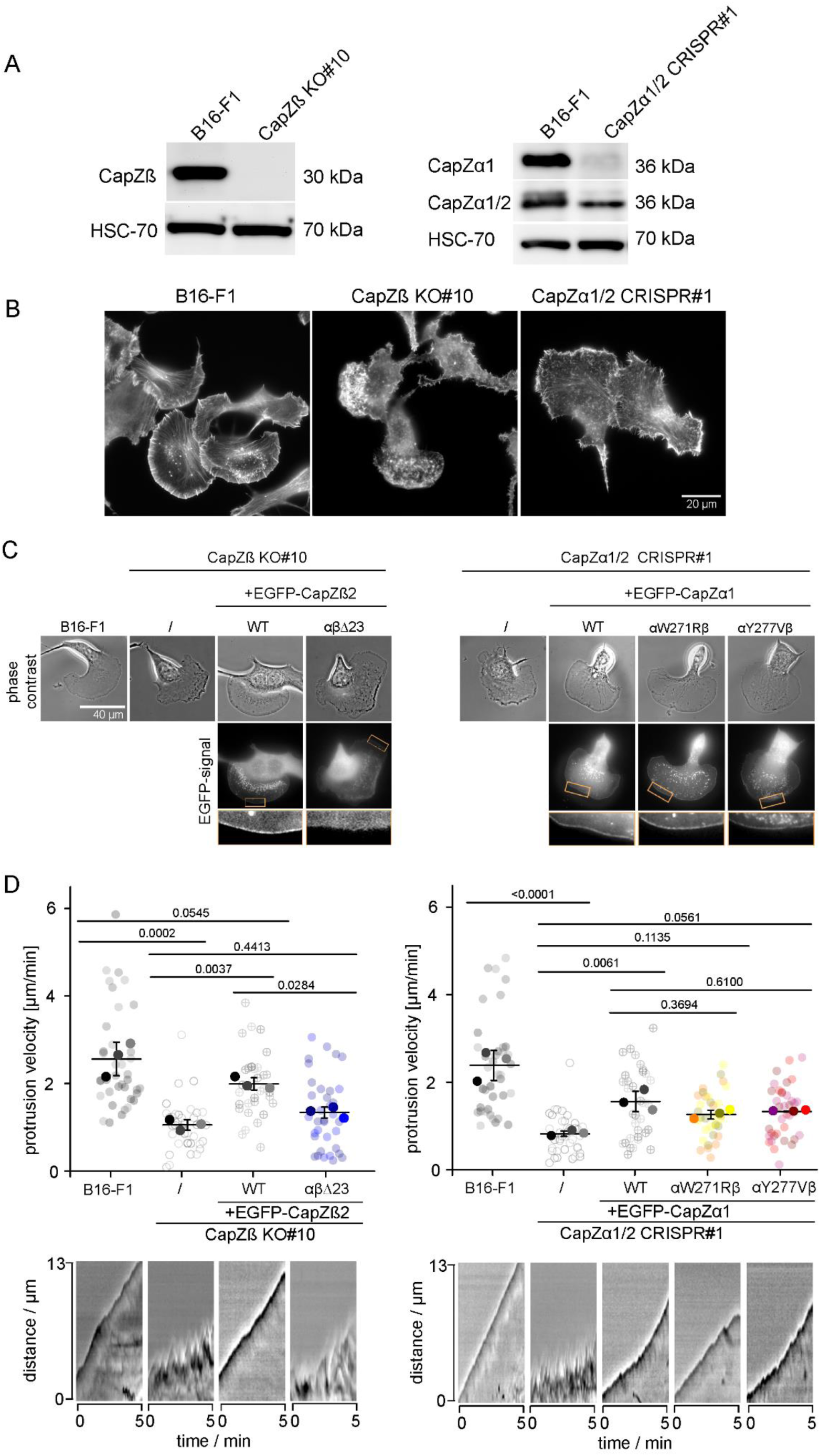
The β tentacle is essential for CP localization to lamellipodial actin networks and efficient leading edge protrusion. (A) Lysates of B16-F1 control cells, cells with CapZβ KO (clone #10, left) and CapZα1/2 CRISPR/Cas9-treated cells (clone #1, right) subjected to western blotting using CapZβ (left) or CapZα1 and CapZα1/2 (right) antibodies. (B) Cell morphologies of B16-F1 *versus* CapZβ KO and CapZα1/2 CRISPR cells stained for the actin cytoskeleton with phalloidin. (C) Representative phase contrast and fluorescence live cell imaging frames of B16-F1 control cells, CapZβ KO and CapZα1/2 CRISPR/Cas9-treated B16-F1 cells with or without expression of EGFP-tagged CapZβ2 (left) or CapZα1 (right) variants, as indicated. Boxed regions (orange) highlight insets at the bottom to reveal localisation patterns of respective, EGFP-tagged rescue constructs. (D) Quantification of protrusion velocity (with representative kymographs at the bottom) of B16-F1 control cells, CapZβ KO (clone 10, left) or CapZα1/2 CRISPR clone 1 (right) cells and the latter expressing CapZβ2 or CapZα1 variants as indicated. P-values were derived from ANOVA Tukey tests.

In agreement with prior knockdown/knockout studies (Hug et al., 1995; Iwasa and Mullins, 2007; Mejillano et al., 2004), functional interference with either CP subunit caused consistent defects in leading edge dynamics reflected by significantly altered morphologies at the peripheries of these cells compared to wild type (Figure 6B, C). These phenotypes coincided with strongly compromised protrusion velocities in each genotype (by app. 70%), which could be largely rescued by EGFP-tagged wildtype variants of respective, disrupted gene (Figure 6C, D). Inefficient protrusion observed in CRISPR/Cas9-treated clones derived from their leading edge membranes undergoing rapid fluctuations, strongly contrasted by the smooth, homogeneous protrusion seen upon rescue with wildtype CPβ and –α as well as the kinetics-mimicking mutants of the latter (αW271Rβ or αY277Vβ, Figure 6D bottom). Furthermore, the restoration of protrusion velocity was similarly efficient with the latter CPα-mutants as seen with wild type CPα (Figure 6D, right panel). As only exception, expression of CPβ lacking its tentacle region in the CPβ-deficient clone was significantly less potent in its rescue efficiency compared to wildtype CPβ with a protrusion efficiency not statistically different from untransfected KO cells (Figure 6D, left panel). The functional difference between CP lacking its β tentacle (αβΔ23) and the corresponding kinetics-mimicking mutants (αW271Rβ or αY277Vβ) also correlated with their accumulation in restored protrusions, since all EGFP-tagged constructs accumulated at protruding cell edges except for αβΔ23 (Figure 6C). Taken together, these data show that the CP β tentacle serves an important function in regulating lamellipodial actin dynamics secondary to simply contributing to high-affinity binding to the barbed ends of actin filaments.

## Discussion

Combining structural and cell biology with in vitro reconstitution, we have elucidated a mechanism that links the two central antagonistic biochemical reactions –capping and nucleation- in the assembly of branched actin networks. Our results identify the WH2 domain of the NPF as the central coupling element that can assume distinct states (Figure 7): 1) Bound to an actin monomer ready to activate the Arp2/3 complex and 2) tethered to a filament barbed end in a nucleation-inactive configuration. We propose that the latter interaction provides negative feedback to Arp2/3 activity and refer to this mechanism as barbed end interference. Importantly, both free, polymerizing barbed ends and capped ends unmasked by the CP β tentacle deletion are capable of sequestering WH2 domains. In the absence of the CP β tentacle, WH2 domains cannot discriminate between capped and free barbed filament ends, effectively creating an additional third state that tethers and inactivates the NPF (Figure 7). We propose that this results in excessive NPF WH2 sequestration and strong inhibition of Arp2/3-dependent branching in the context of the motile network.

**Figure 7.**
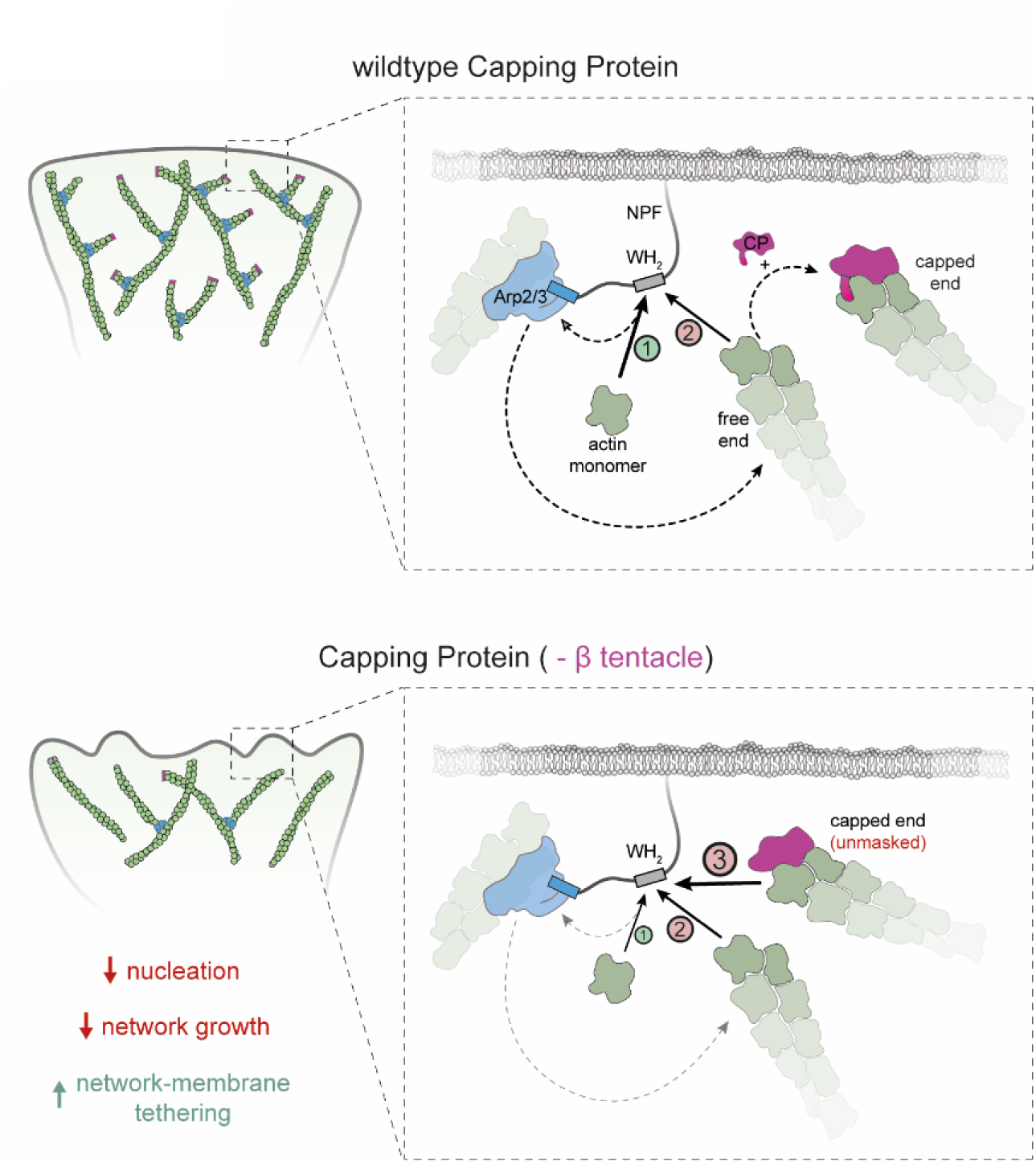
Scheme of the barbed end interference model. Top panel: Actin monomers compete with free barbed ends for NPF WH2 binding. Binding of the former generates the nucleation-competent NPF state (1), whereas binding of the latter sequesters the NPF in an inactive configuration (2). Binding of wildtype capping protein masks the terminal subunits preventing NPF interaction, thereby indirectly stimulating nucleation. Bottom panel: Deletion of the β tentacle unmasks the terminal actin protomer, allowing capped ends to tether additional NPF proteins in an inactive state (3). This strengthens the tethering between actin network and the membrane and decreases the nucleation rate and network growth velocity.

Barbed end interference clearly differs from the previously proposed monomer gating hypothesis (Akin and Mullins, 2008). Monomer gating assumes a kinetic competition between free barbed ends and the Arp2/3 complex for a limiting “flux” of monomeric actin transiently bound to the nucleation promoting factor. Barbed end interference on the other hand simply reflects a direct interaction between NPFs and filament barbed ends. Our results strongly argue for the latter, because a significant fraction of NPFs bind to actin filament ends even when network assembly is kinetically arrested i.e. in the total absence of monomer flux (Figure 5, (Co et al., 2007)).

WH2 domains of WAVE-family proteins bind actin monomers more tightly (K_D_<1 µM) than filament barbed ends (K_D_>10 µM) (Bieling et al., 2018; Chereau et al., 2005; Marchand et al., 2001). How can barbed ends –either free and polymerizing or capped and “unmasked” by the loss of the CP β tentacle–compete with monomers for NPF binding given this large difference in affinity? Several factors might strongly tip the balance towards end binding. First, the density of polymerizing ends at the active front of barbed networks is extremely high (500-2000/μm^2^, (Akin and Mullins, 2008; Bieling et al., 2016; Vinzenz et al., 2012)). Second, in the context of a growing actin network, free ends are confined to the immediate proximity of the NPF, which is attached to the boundary they push against (Bieling et al., 2016; Vinzenz et al., 2012). This type of dimensionality reduction might greatly favor association. Third, bare actin monomers unlikely exist at levels significantly above their critical concentration under physiological conditions (c_c_<1 µM). Indeed, the majority of soluble actin is bound to profilin in living cells (Funk et al., 2019; Kaiser et al., 1999), which itself competes with and reduces the monomer occupancy of NPF WH2 domains (Bieling et al., 2018). The exact contribution of each of these mechanisms to barbed end interference remains to be tested.

In addition to resolving the stimulating effect of CP on Arp2/3-dependent nucleation, barbed end interference explains a multitude of observations otherwise hard to rationalize. Most importantly, it explains why Arp2/3 activation by WASP-family proteins requires the transfer of an actin monomer bound to the WH2 domain. The recent discovery of a second class of Arp2/3 activators devoid of an actin monomer binding site (Balzer et al., 2018; Helgeson and Nolen, 2013) demonstrates that monomer delivery is not a mandatory step in nucleation (Shaaban et al., 2020). Instead, we propose that the NPF WH2 domain evolved as an important, conserved feedback element that keeps autocatalytic branching at bay (Mullins et al., 2018). Barbed end interference also provides an explanation for the universal conservation of the CP β tentacle. Previous genetic and biochemical work indicated that this secondary end-binding site might be dispensable for capping and contributes little to the termination of filament growth as such (Kim et al., 2004; Wear et al., 2003). We show here that it nonetheless serves a critical function in branched network assembly, which is to prevent the capped filament end from WH2 binding. All of these points highlight that the proteins constituting the core branched network motor –Arp2/3, CP and NPFs-are more intimately linked than previously appreciated and have evolved as a functional unit. This illustrates the necessity to study them not only individually as done traditionally, but rather collectively under realistic boundary conditions when constructing a force-generating network.

Several actin-binding proteins unrelated to CP such as gelsolin-family members or epidermal growth factor receptor kinase substrate 8 (EPS8) possess barbed end capping activity (Cooper and Sept, 2008; Edwards et al., 2014). However, these proteins are less conserved within the eukaryotic domain and display a more limited expression across distinct cell types and tissues. Whether their inability to compensate for the loss of CP is the result of a distinct mode of binding at the filament barbed end that does not result in competition with NPF WH2 domains remains to be investigated. Competition for overlapping binding sites at the exposed barbed end face of terminal subunits likely represents a general mechanism by which end-binding proteins reciprocally control their activity. For instance, twinfilin likely removes CP from the barbed end by competition with the tentacles of both CP subunits for actin binding (Hakala et al., 2021; Mwangangi et al., 2021). Likewise, formins can interact with capping protein to accelerate barbed end uncapping (Bombardier et al., 2015; Shekhar et al., 2015). Although structural details are not known, it is likely that this also occurs by competition for partially overlapping sites involving the exposed central barbed end cleft. Finally, we have recently shown that formins accelerate the release of profilin from the barbed end they processively elongate (Funk et al., 2019). Whether this is the result of direct competition or due to allosteric communication remains to be resolved. The structural approach established here provides a powerful strategy to study how all of these and other unrelated end-binding proteins target actin filament ends.

## Acknowledgements

We thank Philippe Bastiaens for continuous support, useful discussions and help in shaping the manuscript. We thank Natalie Petek and Dyche Mullins for Acanthamoeba castellani actin, Dorothy Schafer for kindly providing EGFP-tagged CPβ2. We are also grateful to members of the Bieling lab and Thomas Surrey for comments on the manuscript.

## Notes

**Abbreviations used in this work**
CP: Capping Protein
Arp2/3: Actin Related Protein 2/3 complex
NPF: Nucleation Promoting Factor
PA: Profilin-Actin

## Supplemental Figures

**Supplemental Figure 1:**
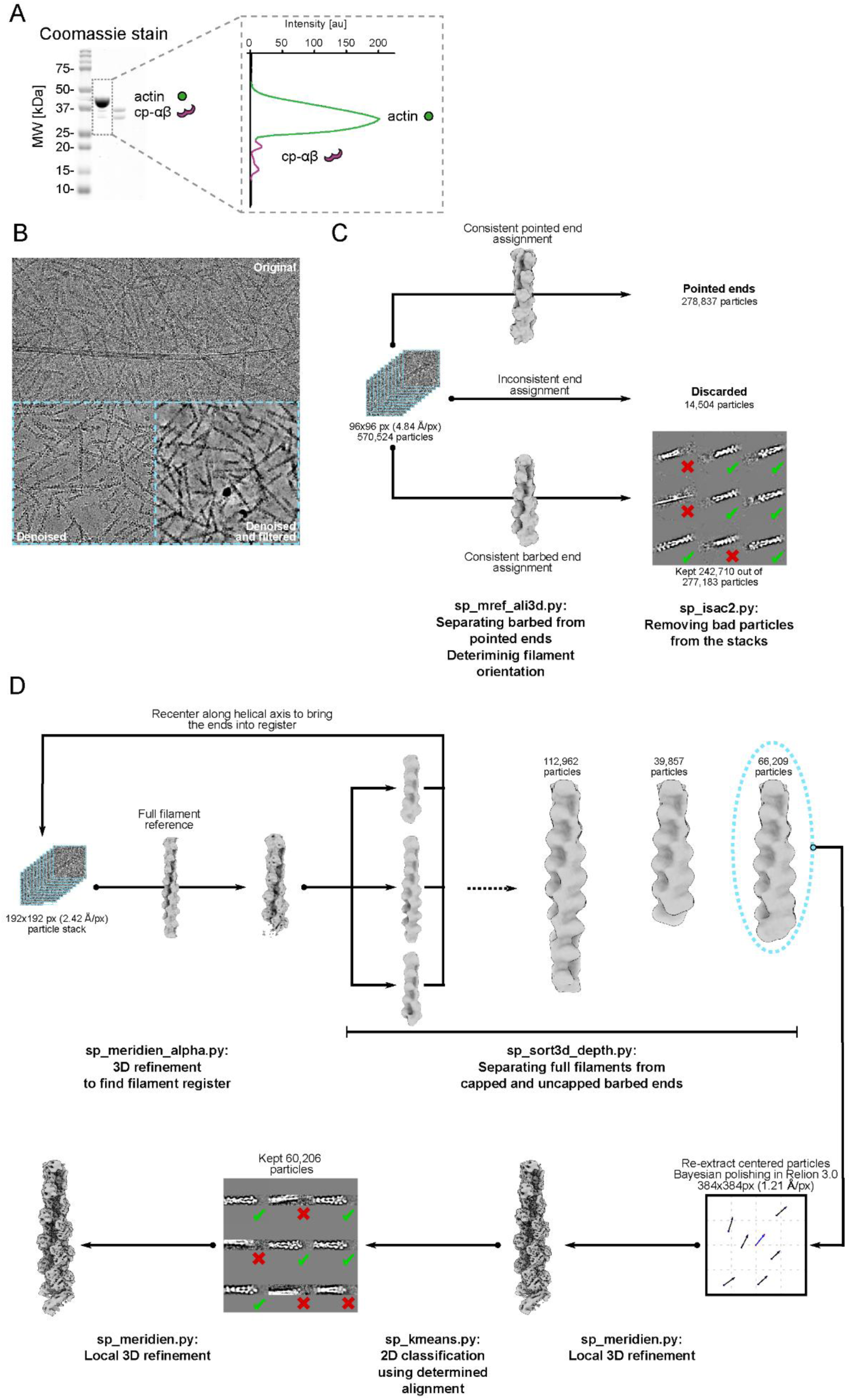
Structure determination of CP-bound filament barbed ends. (A) Protein gel from size-eclusion chromatography. Left: Original Coomassie-stain with visible bands representing actin and α, β-capping protein (see Figure 1 B). Right: Intensity plot of representative protein bands for actin (green) and capping protein (magenta). (B) – (D) Image processing strategy used to obtain the Cryo-EM map of capped filaments. (B) Representative micrograph imaged at ∼ −1.5 µm defocus. The contrast of the images was enhanced with the neural-network denoiser janni, and subsequently with a ctf-correction filter. This was necessary to train a robust crYOLO model for autopicking. (C) Pointed and barbed ends were separated using 3D multi-reference alignment. Only those particles that were assigned consistently in two independent runs were labeled as either pointed or barbed ends. (D) 3D-refinement strategy. 3D-refinement runs were started with a full filament as reference. Initially, the filament ends were located in different monomers along the filament. Using focused 3D classification, we separated them, brought them into register, and used them for a new round of 3D refinement. After the final classification, a single population of capped ends could be identified. Those were subjected to Bayesian polishing (Scheres, 2012), and further refined. A final step of k-means classification using the previously determined euler angles served to remove the last remaining back picks. This was followed by a final local refinement to produce the final map.

**Supplemental Figure 2:**
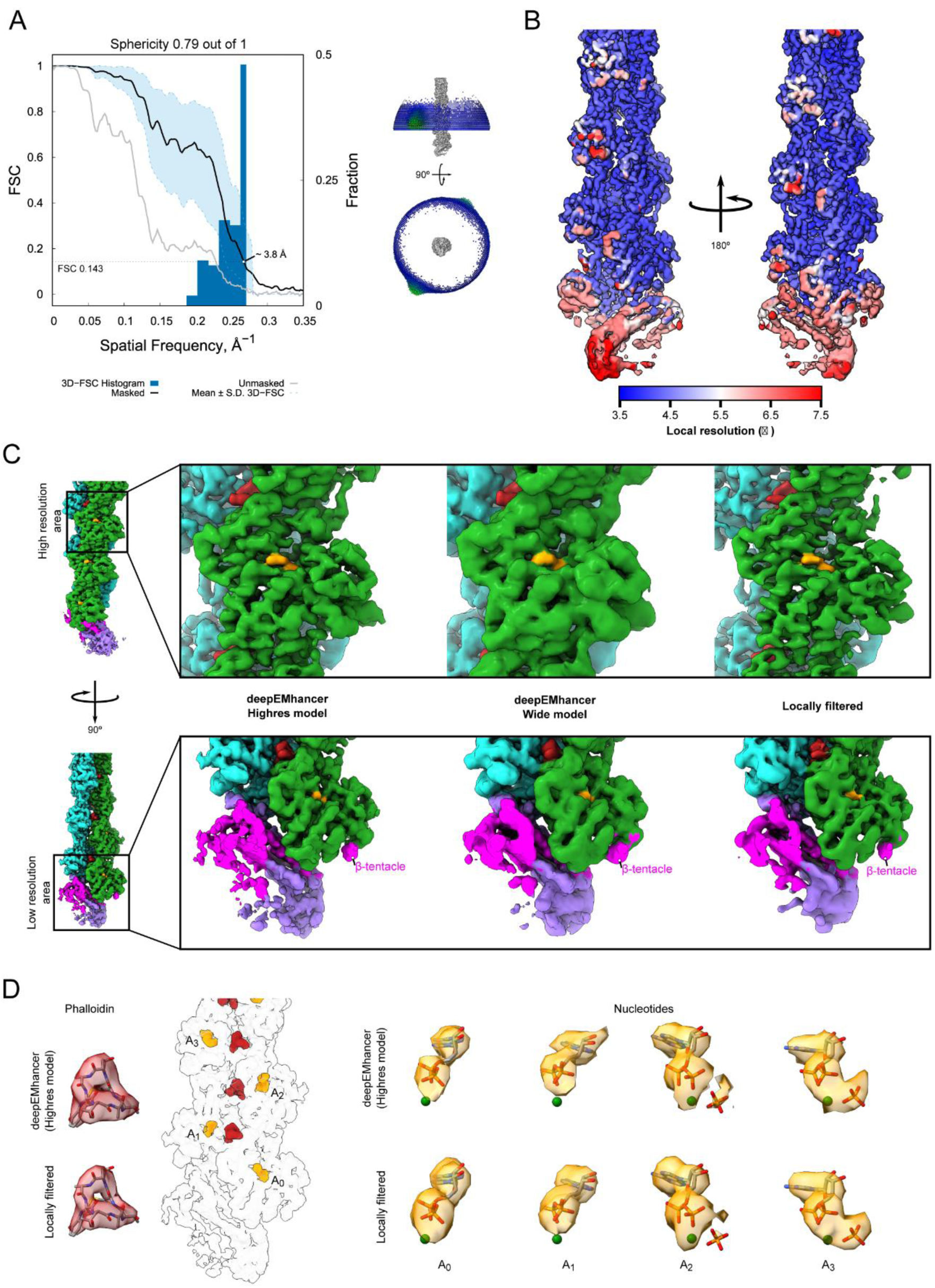
Resolution estimation for the Cryo-EM map. (A) Resolution estimation using Fourier shell correlations. The solid black and gray lines show the overall FSC curves for the map with or without applying a soft mask. The dashed light blue lines and the region enclosed by them represent the average ± one standard deviation of the directional masked FSC curves calculated by 3D-FSC (Tan et al., 2017). A moderate preferential orientation is visible in these curves as well as in the distribution of Euler angles shown on the right of the plot. (B) Local resolution estimated using the false discovery rate procedure implemented in SPOC (Beckers and Sachse, 2020). The color of the map, post-processed by deepEMhancer, goes from low resolution in red to highest resolution in blue. (C) Appearance of the map in regions of high or low local resolution. For each region, maps post-processed with the highres or wide models of deepEMhancer or locally filtered with SPOC are presented. Colors are is in figure 2. (D) Small molecule density found in the Cryo-EM map. The map in the center highlights the position of the small molecules found along the filament. (left) Representative Phalloidin density. (right) Density at the nucleotide-binding site of the first 4 actin monomers. Clear density for ADP can be seen in all of them. The two terminal protomers lack additional density for P_i_, which is present in all remaining active sites. The panels include the density coming from maps post-processed with deepEMhancer or locally filtered.

**Supplemental Figure 3:**
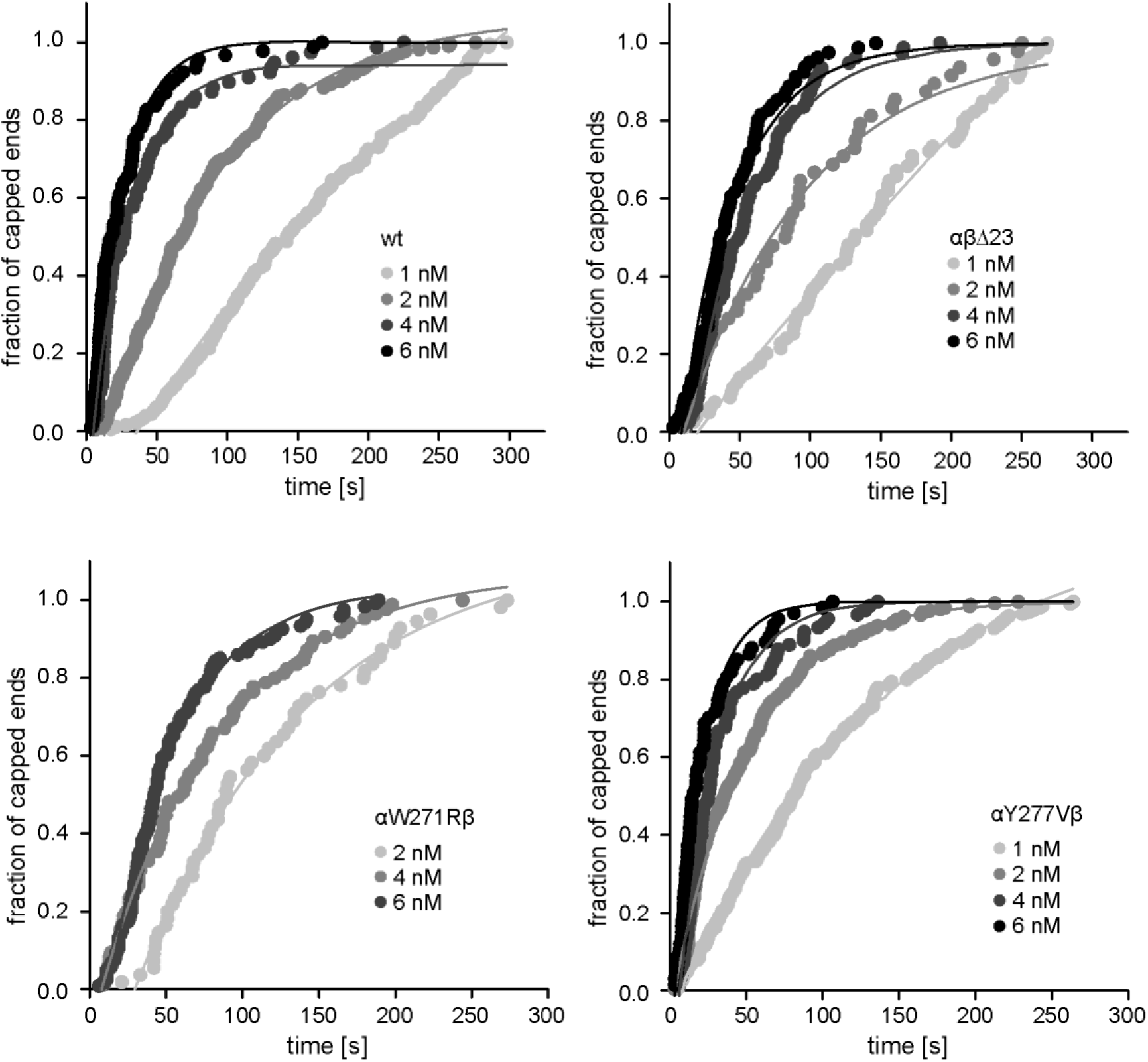
Determination of the barbed end filament association rate constants for capping protein wt and mutants. Rate of actin filament capping in the presence of 2 μM Mg-ATP-actin (10 nM Alexa488-Lifeact), 2 μM profilin1, 1–6 nM capping protein (wt or mutant, N ≥ 80 filaments tracked). The observed reaction rates (k_obs_) (plotted in Figure 3 C) were derived from fits to a mono-exponential growth function (see Materials and methods).

**Supplemental Figure 4:**
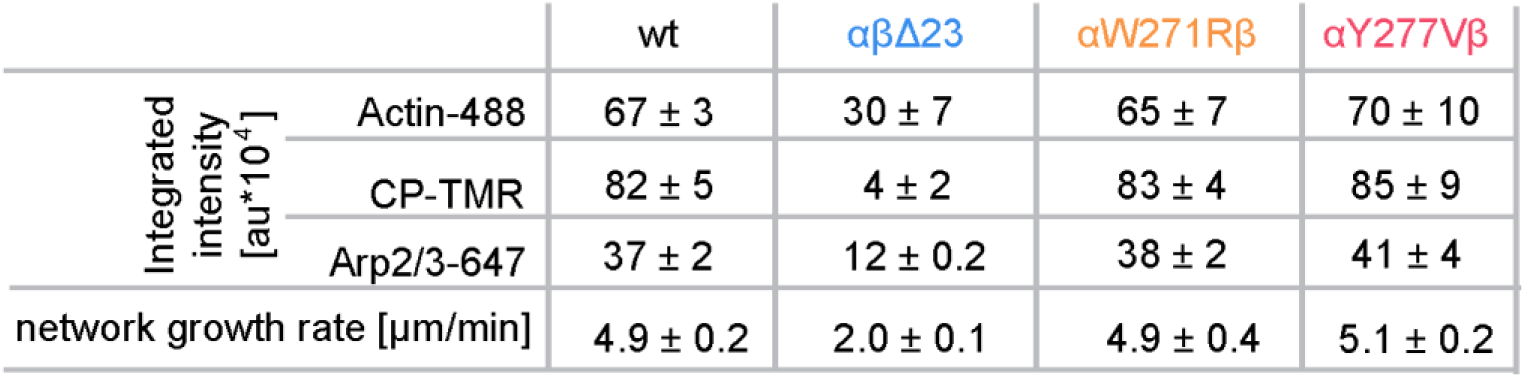
Summary table. This table summarizes the mean integrated intensities and the network growth rates (μm/min) of the actin, the capping protein and the Arp2/3 fluorescence signals for dendritic actin networks grown in the presence of wt or mutant capping protein, error = SD.

**Supplemental Figure 5:**
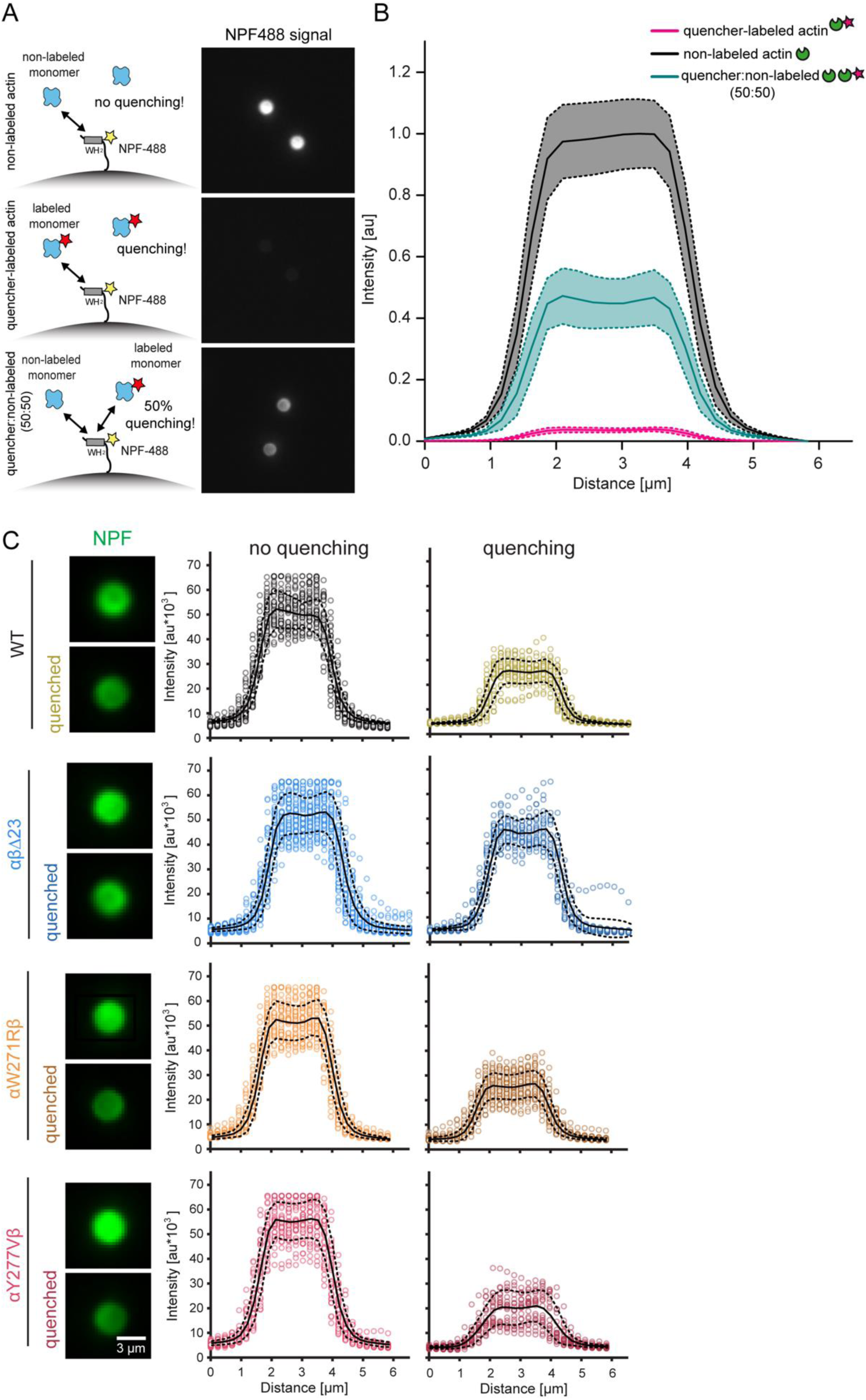
Control experiments for NPF-fluorescence quenching by quencher-labeled actin monomers in bead motility assays. (A) Left: Scheme of experimental setup. Biotinylated, Alexa-488-labeled NPF molecules were bound to the surface of microspheres by a streptavidin linker. The NPF-coated microspheres were incubated with unlabeled (top), quencher-labeled (middle) or a mix of labeled and unlabeled (50:50, bottom) actin monomers, under non-polymerizing conditions, as indicated. Right: Representative images of the NPF-Alexa488 fluorescence signal intensity. (B) Average NPF-fluorescence signal intensities (with error = SD) from NPF-coated microspheres incubated with non-labeled and quencher-labeled actin monomers as indicated in (A). (n ≥ 45 per condition).(C) Left: Representative images from NPF-Alexa488-coated microspheres (see Figure 4 B) acquired in wide field epifluorescence 3 min after arrest before and after quenching with quencher-labeled actin monomers. Networks were grown in presence of wt or mutant capping proteins, as indicated. Right: single point intensities (dots) and average intensities (lines, with error = SD/dotted lines) of the NPF-fluorescent signal intensity on the microsphere surface in presence (quenching) and absence (no quenching) of quencher-labeled actin monomers (N ≥ 25 per CP-type and condition).

## Materials and Methods

### Key Resource Table

| **Key Resource Table** |  |  |  |  |
| --- | --- | --- | --- | --- |
| **Reagent type (species) or resource** | **Designation** | **Source or reference** | **Identifiers** | **Additional Information** |
| Strain, strain background (*Escherichia coli*) | BL21 Rosetta | Novagen | Cat# 70954 | Chemically competent cells |
| Strain, strain background (*Escherichia coli*) | BL21 Star pRARE | EMBL Protein Expression Facility |  | Chemically competent cells |
| Cell line (*mus musculus*) | B16F-1 | ATCC | Cat# CRL-6323, RRID:CVCL_0158 | CP quantifications by WB, CP KO and rescue experiments and EGFP-CP localization in vivo |
| Antibody | anti-CAPZA1 (mouse monoclonal) | BD Transduction Laboratories | 612460 | WB (1:5000) |
| Antibody | anti-CAPZB (mouse monoclonal) | Hybridoma bank USA | 3F2.3 | WB (1:1000) |
| Antibody | anti-CAPZA1/2 (rabbit polyclonal) | Arigo Laboratories | Cat# ARG58328 | WB (1:1000) |
| Antibody | anti-mouse IgG (goat polyclonal) | Dianova | 115-035-062 | WB (1:5000) |
| Antibody | anti-rabbit IgG (goat polyclonal) | Dianova | 111-035-045 | WB (1:5000) |
| Antibody | anti-HSC-70 (mouse monoclonal) | Santa Cruz | sc-7298 | WB (1:10000) |
| Transfected construct | EGFP-CapZα1 | This paper | Uniprot: P52907 | transfected construct, can be obtained in the lab of P. Bieling, MPI Dortmund |
| Transfected construct | EGFP-CapZβ2 | Schafer et al., 1998 → PMID: 9864364 | U10407.1 | Construct can be obtained from Dorothy A Schafer |
|  |  |  |  | University of Virginia, Charlottesville, VA, USA |
| Peptide, recombinant protein | Native bovine $\beta$, $\gamma$ - actin | (Funk et al., 2019) | | For cryo structure and biochemical experiments |
| Peptide, recombinant protein | Bovine Arp2/3 | This paper |  | For the generation of actin dendritic networks |
| Peptide, recombinant protein | Human myotrophin | This paper | Uniprot: P58546 | construct can be obtained in the lab of P. Bieling, MPI Dortmund |
| Peptide, recombinant protein | Mouse capping protein α1 | (Bieling et al., 2018) | Uniprot: 47753 | construct can be obtained in the lab of P. Bieling, MPI Dortmund |
| Peptide, recombinant protein | Mouse capping protein β2 | (Bieling et al., 2018) | Uniprot: 47757 | construct can be obtained in the lab of P. Bieling, MPI Dortmund |
| Peptide, recombinant protein | UTRN<sub>261</sub> | (Bieling et al., 2018) | Uniprot: P46939 | construct can be obtained in the lab of P. Bieling, MPI Dortmund |
| Peptide, recombinant protein | lifeact | (Bieling et al., 2018) |  | construct can be obtained in the lab of P. Bieling, MPI Dortmund |
| Peptide, recombinant protein | Human profilin 1 | (Funk et al., 2019) | Uniprot: P07737 | construct can be obtained in the lab of P. Bieling, MPI Dortmund |
| Peptide, recombinant protein | Human WAVE1 WH<sub>2</sub> PVCA | (Bieling et al., 2018) |  | construct can be obtained in the lab of P. Bieling, MPI Dortmund |
| Peptide, recombinant protein | Human N-WASP | (Bieling et al., 2018) |  | construct can be obtained in the lab of P. Bieling, MPI Dortmund |
| Chemical compound, drug | Latrunculin B | Sigma Aldrich | Cat. #: L5288 | For actin arrest |
| Chemical compound, drug | Phalloidin | Sigma Aldrich | Cat. #: P2141 | For actin arrest |
| Chemical compound, drug | EZ-Link MaleimidePEG2-Biotin | Thermo Fisher | Cat. #: A39261 | For NPF labeling |
| Chemical compound, drug | HO-PEG-NH2 and Biotin-CONH-PEG-OC3-H6-CONHS | Rapp Polymere | # 103000–20 and # 133000-25-35 | For glass surface functionalization |

### Protein biochemistry – Purification and labeling

#### Native (β, γ) -actin

Native bovine (β, γ) –actin was purified from bovine thymus tissue according to the methods described previously (Funk et al., 2019). Briefly, (β, γ) –actin was purified from fresh bovine thymus tissue. After lysis of the thymus tissue, the solution was hard spun and filtered to generate a cleared supernatant that was incubated with 10xhis-gelsolinG4-6 fragment to promote the formation of actin:gelsolin G4-6 complexes. Next, the lysate was circulated over a Ni^2+^ superflow column. Actin monomers were eluted and polymerized into filaments. After ultracentrifugation, actin filaments were resuspended in F-buffer (1xKMEI, 1xBufferA) and stored in continuous dialysis at 4 °C. F-buffer containing fresh ATP and TCEP was continuously exchanged every 4 weeks. The actin was depolymerized through dialysis into BufferA (2 mM Tris, 0.2 mM ATP, 0.1 mM CaCl_2_, 0.5 mM TCEP) for 6 days and gelfiltered over a Superdex 200 16/600 column prior to all experiments.

#### Profilin and profilin-actin complexes

Human profilin1 and stoichiometric complexes from profilin and β, γ –actin were purified as described in (Bieling et al., 2018; Funk et al., 2019). Briefly, human profilin1 was expressed as untagged protein in *E. coli* Rosetta cells at 30°C for 4.5 hr. Profilin1 was purified by ammonium sulfate precipitation, followed by ion-exchange (DEAE) and hydroxylapatite (HA) chromatography steps, followed by size exclusion chromatography (Superdex 200 16/600) into storage buffer (20 mM Tris-Cl pH 7.5, 50 mM NaCl, 0.5 mM TCEP). Proteins were snap-frozen in liquid nitrogen upon addition of 20% glycerol to the storage buffer, and stored at −80°C.

To generate profilin-actin complexes, freshly gelfiltered actin monomers were incubated with 1.5x molar excess of profilin1 at 4°C overnight. Next, profilin-actin complex was separated from free profilin by gelfiltration over a Superdex 200 10/300 GL into Buffer A. The complex was concentrated to working concentrations between 100 and 200 μM and stored at 4 °C up to two weeks without inducing nucleation.

#### Atto540Q-quencher labeled actin

For the labeling of actin monomers with Atto540Q-NHS, the complex of human profilin1 and *Acanthamoeba castellani* actin (Hansen et al., 2013) was generated in Buffer A using a 2x molar excess of profilin1. Following gelfiltration into labeling buffer (2 mM Hepes pH 8.0, 0.1 mM CaCl_2_, 0.2 mM ATP, 0.5 mM TCEP), the profilin-actin complex was labeled at its reactive lysine residues by incubating with a 3x-fold excess of Atto540Q-NHS for 40 min at 23 °C. The reaction was terminated by adding Tris-Cl (pH 8.0) to a final concentration of 2 mM to the reaction mix. After 10 min incubation, soluble dye and protein were separated from solid particles by ultracentrifugation for 10 min at 300.000xg. Next, polymerization of actin was initiated by adding KMEI-buffer (10 mM imidazole pH 7.0, 50 mM KCl, 1.5 mM MgCl_2_, 1 mM EGTA) and 1 % of pre-polymerized, unlabeled, freshly-sheared actin filaments to the labeling mix (Bieling et al., 2016). After 1 hr of polymerization at 23 °C, filaments were separated from excess soluble dye, non-polymerizable actin and profilin by ultracentrifugation for 10 min at 300.000xg. Actin filaments were stored in F-buffer dialysis at 4 °C and dialyzed into Buffer A (for 6 days) prior to experiment. The degree of labeling (73–87 %) was determined by absorbance at 280 nm and 543 nm.

#### Myotrophin / V1

Human myotrophin was expressed in *E. coli* BL21 Rosetta cells from a pETM11 (Bieling et al., 2018) vector containing a N-terminal 10xhis tag followed by a TEV-cleavage site. After protein expression for 16 hrs at 18 °C, cells were lysed (50 mM KP_i_ pH 7.3, 400 mM NaCl, 10 mM imidazole, 1 mM β-mercaptoethanol, 1 mM PMSF, DNaseI). The lysate was hard spun and purified by IMAC over a 5 ml HiTrap Chelating column. After gradient elution (50 mM KP_i_ pH 7.3, 400 mM NaCl, 10 mM imidazole, 1 mM β-mercaptoethanol), the 10xhis-tag was cleaved by TEV protease overnight. After cleavage, the protein was desalted into lysis buffer and filtered over a HiTrap Chelating column. The flow through was gelfiltered over a Superdex 200 column into storage buffer (20 mM Hepes pH 7.5, 50 mM KCl, 0.5 mM TCEP) and snap-frozen in liquid nitrogen supplemented with 20 % glycerol. For long-term storage, the protein was stored at −80 °C.

#### Capping protein

Capping protein mutants were either generated via PCR amplification and Gibson cloning (Gibson, 2011) (βΔ23) or via site-directed mutagenesis (αW271R, αY277V, EGFP-CapZb2). Murine heterodimeric capping protein (α1 in pETM20, β2 in pETM33 (Bieling et al., 2018)) were co-expressed in *E. coli* BL21 Rosetta cells. Following protein expression, cells were lysed (50 mM K_2_PO_4_ pH 7.3, 400 mM NaCl, 5 mM imidazole, 1 mM EDTA, 2 mM PMSF, 0.75 mM β-mercaptoethanol, 15 μg/ml benzamidine, DNaseI) and purified by IMAC over a 5 ml HiTrap Chelating column. After gradient elution (50 mM K_2_PO_4_ pH 7.3, 400 mM NaCl, 400 mM imidazole, 1 mM EDTA, 2 mM PMSF, 0.75 mM β-mercaptoethanol), the his-tag was cleaved by TEV/Precision proteases overnight. Following cleavage, the protein was desalted into low salt MonoQ buffer (10 mM Tris-Cl pH 8.0, 5mM KCl, 1 mM EDTA, 2 mM PMSF, 1 mM DTT) and again filtered over a HiTrap Chelating column followed by a MonoQ run. After protein elution (10 mM Tris-Cl pH 8.0, 1 M KCl, 1 mM EDTA, 2 mM PMSF, 1 mM DTT) the protein was gelfiltered over a Superdex 200 column (10 mM Tris-Cl pH 7.5, 50 mM KCl, 0.5 mM TCEP, 20 % glycerol), concentrated and either stored in −80 °C or directly transferred into a SNAP-labeling reaction.

To generate fluorescently-labeled capping protein, an N-terminal SNAP-tag (Keppler et al., 2003) was fused to the beta subunit. For N-terminal SNAP-labeling, a 3x molar excess of SNAP Cell TMR-star was mixed with the protein and incubated for 4 hrs at 16 °C following an overnight incubation on ice. Next, the protein was gelfiltered over a Superose 6 10/300 GL column into storage buffer. The degree of labeling (50–70%) was determined by absorbance at 280 nm and 554 nm. The addition of N-terminal SNAP-tags did not affect the activity of capping protein as measured in single molecule/filament TIRF-M assays and bead motility experiments.

#### Arp2/3 complex

Native bovine Arp2/3 complex was purified from fresh bovine thymus by ammonium sulfate precipitation and ion exchange chromatography (DEAE, Source Q and Source S) followed by gelfiltration over a Superdex 200 as previously described (Bieling et al., 2016; Doolittle et al., 2013). The protein was either directly snap frozen with 20 % glycerol and stored at −80 °C or transferred into a labeling reaction.

Arp2/3 was fluorescently labeled by the addition of 5x-fold molar excess of maleimide-Alexa647 dye conjugate. After incubation on ice for 2 hrs, the reaction was terminated by adding 2 mM DTT for 30 min. Next, the complex was loaded onto an N-WASP-VCA column (5 mM Tris-Cl pH 8.0, 50 mM NaCl, 0.5 mM MgCl_2_, 0.1 mM ATP, 2 mM DTT) and gradient-eluted over 100 column volumes (10 mM Tris-Cl pH 8.0, 1 M NaCl, 0.5 mM MgCl_2_, 0.1 mM ATP, 2 mM DTT). Fractions containing the full complex were gelfiltered (5 mM Hepes pH 7.5, 50 mM KCl, 0.5 mM EGTA, 0.5 mM MgCl_2_, 0.1 mM ATP, 0.2 mM TCEP) over a Superose 6 column, concentrated, snap frozen with 20 % glycerol and stored at −80 °C.

#### Preparation of an N-WASP-loaded column for Arp2/3 complex labeling

Before immobilizing the N-WASP-VCA domain onto a 1 ml HiTrap NHS activated HP column, 10 mg of the protein were desalted into coupling buffer (50 mM Hepes pH 8.0, 500 mM NaCl, 0.5 mM DTT). The column was pretreated according to the recommendations of the manufacturer (GE healthcare). After all protein had bound to the resin, the column was equilibrated with wash buffer (5 mM Tris-Cl pH 8.0, 50 mM NaCl, 0.5 mM MgCl_2_, 0.1 mM ATP, 2 mM DTT).

#### N-WASP VCA

Human N-WASP VCA was expressed with an N-terminal 10xhis-GST-tag using *E. coli* BL21 Star pRARE. After expression for 16 hrs at 18 °C, cells were lysed (100 mM KP_i_ pH 7.2, 1 M NaCl, 0.5 mM β-mercaptoethanol, 5 mM imidazole, 1 mM PMSF, 15 μg/ml benzamidine, DNaseI) and protein was purified by IMAC over a 5 ml HiTrap Chelating column. After gradient elution, tags were removed by TEV protease cleavage overnight. After desalting the protein back into lysis/wash buffer, non-cleaved protein and free 10xhis tag was removed by passing the protein over a 5 ml HiTrap Chelating column. Next, the protein was loaded onto a MonoQ column (20 mM Tris-Cl pH 8.0, 50 mM NaCl, 1 mM DTT) and gradient eluted over 30 column volumes (20 mM Tris-Cl pH 8.0, 1 M NaCl, 1 mM DTT) followed by gelfiltration over a Superdex 200 column (20 mM Tris-Cl pH 8.0, 150 mM NaCl, 1 mM DTT). The protein was snap-frozen in liquid nitrogen with 20 % glycerol and stored at −80 °C.

#### WAVE WH2

Human WAVE1 PVCA WH2 was expressed with an N-terminal 10xhis-ztag followed by penta-glycine and a C-terminal KCK motif in *E. coli* BL21 Star pRARE. After expression for 16 hrs at 18 °C, cells were lysed (50 mM Tris-Cl pH 8.0, 150 mM KCl, 0.5 mM β-mercaptoethanol, 1 mM PMSF, 15 μg/ml benzamidine, DNaseI), and protein was purified by IMAC over a 5 ml HiTrap Chelating column. After gradient elution (50 mM Tris-Cl pH 8.0, 150 mM KCl, 0.5 mM β-mercaptoethanol, 300 mM imidazole), tags were removed by TEV protease cleavage overnight. After desalting the protein back into lysis/wash buffer, non-cleaved protein and free 10xhis tag was removed by passing the protein again over a 5 ml HiTrap Chelating column. Next, the protein was gelfiltered over a Superdex 75 column (10 mM Tris-Cl pH 7.5, 50 mM KCl, 0.5 mM TCEP). The protein was snap-frozen in liquid nitrogen with 20 % glycerol and stored at −80 °C.

#### NPF-WAVE-biotinylation by maleimide chemistry

NPF (WAVE(PVCA) WH2) molecules were labeled with EZ-Link maleimide-PEG_4_-Biotin (Thermo Fisher Scientific) at the C-terminally located KCK, as recommended by the manufacturer. Briefly, NPF-KCK was desalted into reaction buffer (5 mM Hepes pH 7.5, 150 mM NaCl, 0.5 mM TCEP), mixed with 10x molar excess of EZ-Link maleimide-PEG_4_-Biotin and incubated for 2 hrs on ice. After quenching the reaction with 1 mM DTT, the protein was gelfiltered over a Superdex 75 10/300 GL into storage buffer and directly transferred into another labeling reaction via sortagging.

#### NPF-PEG-Biotin-Alexa 488/or mCherry by sortagging

Following biotinylation, the NPF was transferred into a second labeling reaction by sortagging. The NPF was labeled with a LPETGG conjugated Alexa-488/or mCherry dye at the N-terminus as outlined in (Antos et al., 2017). Briefly, the NPF-protein was mixed with sortase (in a 3:1 molar ratio), Alexa488-LPETGG conjugate in 4 molar excess, 6 mM CaCl_2_, 7.78 mM Tris pH 8.0, 150 mM KCl, 0.5 mM TCEP. After 18 hrs of incubation at 18 C°, the protein was separated from sortase and free dye by gelfiltration over a S200 10 300 GL column and snap-frozen in liquid nitrogen. The degree of labeling was determined at 20/496 nm.

### Biochemical assays

#### Buffers

All experiments were carried out in a common final assay buffer of the following composition if not stated otherwise: 20 mM Hepes pH 7.0, 100 mM KCl, 1.5 mM MgCl2, 1 mM EGTA, 20 mM β-mercaptoethanol, 0.1 mg/ml β-casein, 1 mM ATP. This buffer has a molar ionic strength of 0.133 M, which is close to the physiological ionic strength found in the literature (between 0.1 and 0.2 M).

#### Functionalization of glass coverslip surfaces and protein immobilization

For TIRF-M experiments, microscopy counter slides were passivated with PLL-PEG and coverslips (22×22 mm, 1.5 H, Marienfeld-Superior) functionalized according to (Bieling et al., 2016; Funk et al., 2019). Reaction flow chambers were blocked with a Pluronic block solution (0.1 mg/ml κ-casein, 1 % Pluronic F-127, 1 mM TCEP, 1xKMEI) followed by 2 washing steps with 40 μl each (0.5 mM ATP, 1 mM TCEP, 1xKMEI, 0.1 mg/ml β-casein). Next, the channels were incubated with 75 nM streptavidin for 4 min following a washing step and incubation of 90 nM biotin-phalloidin (Funk et al., 2019; Hansen and Mullins, 2010; Pollard, 1982) for 4 min.

To visualize actin filament elongation in TIRF-M, a modified protocol outlined in (Hansen and Mullins, 2010) and (Kuhn and Pollard, 2005) was performed. Briefly, 9 μl of a 4,44x μM profilin-actin solution was incubated with 1 μl of 10x ME (0.5 mM MgCl_2_, 2 mM EGTA) for 1 min at RT following the addition of 4 μl oxygen scavenging mix (1.25 mg/ml glucose-oxidase, 0.2 mg/ml catalase, 400 mM glucose) (Aitken et al., 2008; Bieling et al., 2010; Rasnik et al., 2006). The Mg-ATP-profilin-actin was then mixed with 26 μl reaction buffer containing TIRF-buffer (20 mM Hepes pH 7.0, 100 mM KCl, 1.5 mM MgCl_2_, 1 mM EGTA, 20 mM β-mercaptoethanol, 0.1 mg/ml β-casein, 0.2% methylcellulose (cP400, M0262, Sigma-Aldrich), 1 mM ATP, 2 mM Trolox) and additives including 10 nM Cy5-UTRN_261_ and others, which are described in the specific methods and results sections. All TIRF-single filament elongation experiments were performed using profilin-actin as polymerizable substrate unless otherwise indicated in the corresponding results sections and figure legends.

### Measuring capping protein association and dissociation kinetics for the filament barbed end

Experiments were essentially performed as described in the previous section with the following modifications:

#### K_on_ measurements of capping protein for filament barbed ends by TIRF-microscopy

To determine the association rate constant for capping protein (wt and affinity mutants) for the actin filament barbed end, experiments were conducted similar to protocols outlined in (Hansen and Mullins, 2010; Kuhn and Pollard, 2005) with modifications. Briefly, small pre-polymerized phalloidin stabilized actin seeds were immobilized to the glass surface of a TIRF chamber. Increasing capping protein (TMR-labeled or non-labeled CP) concentrations (0 – 6 nM, wt or mutant as indicated) were combined with 2 μM Mg-ATP actin monomers and 2 μM profilin1 in TIRF reaction buffer and flowed into the chamber. Elapsed time between mixing and imaging was ∼40 s. The time courses of capping protein binding to the filament barbed ends (termination of barbed end polymerization) was recorded in single molecule or single filament TIRF time-lapse experiments. Only filaments that were visualized in the first frame were followed throughout the experiment. Filaments that were immobilized from solution during the experiment or new filaments that originated from spontaneous nucleation were excluded from the analysis. Therefore, only filament barbed ends that were initially free were scored. The experiment was recorded over a time period of 5 min in 1 s intervals. ≥ 80 filaments were measured for each capping protein concentration from ≥ 3 slides.

#### K_off_ measurements of capping protein from filament barbed ends by TIRF-microscopy

To determine the dissociation rate constant for capping protein (wt and affinity mutants) from the actin filament barbed end, experiments were conducted similarly to previously-described protocols (Hansen and Mullins, 2010), with modifications. Briefly, pre-polymerized actin filaments were capped by mixing them with excess of capping protein (200 nM non-labeled CP, wt or mutant as indicated). After immobilizing the capped actin filaments in a flow chamber, capping protein was washed out by flushing the chamber with 2x 40 μl of wash buffer. Following wash, 2 μM of profilin-actin and 20 nM myotrophin/V1 (in TIRF buffer) were flushed into the chamber and the time courses of filament re-growth (capping protein dissociation) recorded in single filament TIRF time-lapse experiments. Elapsed time between mixing and imaging was ∼40 s. Only filaments that were visualized in the first frame were followed throughout the experiment. Filaments that were immobilized from solution during the experiment or new filaments that originated from spontaneous nucleation were excluded from the analysis. Therefore, only filament barbed ends that were initially capped were scored in the analysis. The experiment was recorded over a time course of 50 min in 30 s intervals. ≥ 50 filaments were measured for each capping protein version (wt or mutant) from ≥ 3 slides.

#### Actin dendritic network assembly assay

Actin dendritic network assembly assays were carried out using similar protocols as (Akin and Mullins, 2008; Bieling et al., 2016), with modifications. In brief, Streptavidin-coated polystyrene beads (Bangs laboratories, Ø=3 μm) were washed 2 times with NPF dilution buffer (20 mM Hepes pH 7.0, 100 mM KCl, 1 mM EDTA, 1.5 mM MgCl_2_, 0.2 mg/ml β-casein, 5 mM β-mercaptoethanol) at 4 °C and spun at 8000xg for 1 min to remove bead storage solution. After 5 min of sonication, the beads were incubated with biotinylated-NPF (mcherry-WAVE-PWCA and dark cherry-WAVE-PWCA, 1:40) for 30 min at 23 °C followed by 2 washing steps to remove non-specifically absorbed protein. To reduce non-specific protein absorption, glass counter surfaces and glass coverslips (22×22 mm, 1.5 H, Marienfeld-Superior) were plasma-treated and passivated with PLL-PEG. To generate actin dendritic network growth on NPF-coated beads, the following components were combined (motility mix): 5 μM profilin-actin, 100 nM CP (20 % TMR-labeled), 100 nM Arp2/3 (20 % Alexa647-labeled), 15 nM Alexa488-Lifeact, mCherry-NPF-coated beads and reaction buffer (20 mM Hepes pH 7.0, 100 mM KCl, 1.5 mM MgCl_2_, 1 mM EGTA, 20 mM β-mercaptoethanol, 0.1 mg/ml β-casein, 0.2% methylcellulose (cP400, M0262, Sigma-Aldrich), 1 mM ATP). After 4 min (as indicated in the specific sections) at 23 °C, network growth was arrested by adding 15 μM latrunculinB and phalloidin to the mixture. Microscopy chambers were prepared by adding 5 μl of arrested bead motility mix between the glass items. The chambers were sealed with valap (Akin and Mullins, 2008) to prevent evaporation. After the beads had settled down on the glass surface, images were acquired in multicolor.

#### FRET-WH_2_-competition assay to determine the NPF-WH_2_-occupancy on CP decorated actin filament ends

Assays were carried out similarly to the previous section with the following exceptions: Actin dendritic networks were grown from donor-labeled Alexa488-NPF (WAVE(PWCA)), CP and Arp2/3 were unlabeled, and 15 nM Cy5-UTRN_261_ was used as actin filament binding probe. The dendritic actin networks were grown for 5 min at 23 °C following an arrest cocktail containing 5 μM myotrophin/V1, 15 μM Latrunculin B, 15 μM phalloidin, 7.5 μM profilin1 and 7.5 μM quencher-labeled and Latrunculin B-stabilized A540Q-actin monomers (all final concentration). The actin dendritic networks were directly imaged after adding the arrest cocktail. As a negative control, non-labeled wt profilin-actin instead of quencher-labeled actin was added to the arrest cocktail and directly mixed with 488-NPF-coated beads. As a positive control, only quencher-labeled profilin-actin was added to the 488-NPF-coated beads, assuming that all WH_2_ domains are free to interact only with quencher-labeled actin monomers in the absence of a dendritic network.

#### Generation of short, uni-sized actin filaments decorated with capping protein

To generate short (70-160 nm) actin filaments, actin monomers were freshly gelfiltered over a Superdex 200. 40 μl of 40-45 μM actin monomers were sufficient to start the polymerization reaction by addition of 25 μM capping protein. After 2 min incubation at 4 °C, 0.5 mM MgCl_2_ and 0.2 mM EDTA final concentration were added for 1 min at 4 °C, followed by initiation of polymerization with 1xKMEI (10 mM imidazole pH 7.0, 50 mM KCl, 1.5 mM MgCl_2_, 1 mM EGTA) for 2 min at 23 °C. To terminate the reaction, 48 μM phalloidin were added to the reaction mix followed by gelfiltration over a Superdex 200 increase 5/150 GL in 1xKMEI buffer. Short, unisized actin filaments could be collected from the first peak after 1 ml and were directly transferred onto cryo carbon grids for negative stain screening and cryo EM data collection. The freshly gelfiltered samples were stable for at least 4 hrs.

#### Integration of protein intensity on SDS gel after SEC

After SDS-gel run, actin and capping coomassie signal intensities were analyzed using ImageJ. After selecting protein signal areas for each protein, intensities were plotted in a plot profile reflecting the intensity signal across the selected area.

#### Cell culture of B16-F1 cells

B16-F1 cells and derived CapZa1/2 CRISPR#1 and CapZb KO#10 were cultured in DMEM (4.5 g/L glucose; Invitrogen, Germany), supplemented with 10% FCS (Gibco), 2 mM glutamine (Thermo Fisher Scientific) and penicillin (50 Units/mL)/streptomycin (50 µg/mL) (Thermo Fisher Scientific) at 37°C and 7.5% CO_2_. Cells were routinely transfected in 35 mm dishes or 6-well plates, using 500 ng DNA in total and 1 µL JetPrime. After overnight transfection, cells were either plated onto acid-washed, laminin-coated (25 µg/mL) coverslips for phalloidin stainings or in µ-Slide 4 Well Ph+ (Ibidi) chamber slides, coated with 25 µg/ml laminin, for live cell fluorescence microscopy.

#### Generation of CapZα1/2 and CapZb CRISPR/Cas9 clones

Disruption of CPα or –β expression in B16-F1 cells was achieved by targeting *CapZb* or *CapZa1* and *CapZa2* simultaneously using CRISPR/Cas9-mediated genome editing, using targeting/guide sequences CCTCAGCGATCTGATCGACC (for *CapZb* exon 2), GAGTTTAATGAAGTATTCAA (for *CapZa1* exon 2), and CAGAAGGAAGATGGCGGATC (for *CapZa2* exon 1), cloned into plasmid pSpCas9(BB)-2A-Puro (Addgene. ID 48139). Plasmids were transfected into B16-F1 cells, followed by selection of transfected cells for three days using 2.5 µg/ml puromycin (Sigma-Aldrich, Taufkirchen, Germany). Puromycin-resistant cells were then extensively diluted and single cell-derived colonies grown for 5 to 7 days. After colony isolation, cells were expanded and clones screened by Western Blotting using CapZb or CapZa1 and CapZa2-specific antibodies, allowing selection of CRISPR-clones #10 (for *CapZb*) and #1 (for *CapZa1/2*). The genotypes of these clones were examined using TIDE sequencing (Brinkman et al., 2014). In case of CapZb KO#10, all functional alleles could be disrupted. Instead, for CapZa1/2 CRISPR clone #1, sequencing revealed that all *CapZa1* alleles were disrupted, whereas expression from *CapZa2* alleles appeared partially possible from alleles harboring a 9 bp deletion. Additional, subsequent targeting of *CapZa2* alleles on exon 2 failed to eliminate expression from those 9 bp-deletion alleles (not shown).

#### Western blotting

For preparation of whole cell lysates, cells were washed with PBS, lysed using Laemmli-buffer (2% SDS, 10% glycerol, 5% β-mercaptoethanol, 0.05% bromophenol blue, 63 mM Tris-HCl pH 6,8), sonicated to shear genomic DNA and boiled for 5 min at 95°C. Western blotting was carried out using standard techniques, and with antibodies as listed in the Key Resources Table. Chemiluminescence signals were obtained upon incubation with ECL Prime Western Blotting Detection Reagent (GE Healthcare), and recorded with ECL Chemocam imager (Intas, Goettingen, Germany).

#### Phalloidin staining

B16-F1, CapZa1/2 CRISPR#1 and CapZb KO#10 cells were fixed with pre-warmed, 4% paraformaldehyde in PBS, supplemented with 0.25 % glutaraldehyde for 20 min, and permeabilized with 0.05% Triton X-100 in PBS for 30 seconds. The actin cytoskeleton was stained using Alexa 488-conjugated phalloidin (dilution 1:200 in PBS). Samples were mounted using ProLong Gold antifade reagent and imaged using a ×63/1.4NA Plan apochromatic oil objective.

#### Fluorescence microscopy data acquisition

##### TIRF-Microscopy data acquisition

All in vitro experiments were performed at 23°C, unless otherwise specified. Images were acquired using a customized Nikon Ti2 TIRF-Microscope with Nikon perfect focus system. TIRF-images were acquired by a dual EM CCD Andor iXon camera system (Cairn) controlled by NIS-Elements software. Dual color imaging was performed through an Apo TIRF 60x oil DIC N2 objective using a custom multilaser launch system (AcalBFi LC) at 488 nm and 560 nm and a Quad-Notch filter (400-410/488/561/631640). Images were acquired at intervals of 1–30s.

##### Wide-field epifluorescence-microscopy data acquisition

All in vitro experiments were performed at 23°C. Bead motility assays were carried out using a Olympus IX81 wide field epifluorescence microscope with LED illumination (pE4000, CollLED). Image acquisition was performed by a Hamamatsu c9100-13 EMCCD camera controlled by Micromanager 1.4 software (Edelstein et al., 2014). Multi color images were taken through an UPlanS APO 40x or 60x oil objective.

##### Live cell fluorescence microscopy and quantification of protrusion velocity

B16-F1, CapZa1/2 CRISPR#1 and CapZb KO#10 cells, untransfected or transfected with indicated EGFP-tagged CapZ constructs, were plated in µ-Slide 4 Well Ph+ (Ibidi) chamber slides overnight. Next day, the medium was replaced with microscopy medium (F12 HAM HEPES-buffered medium, Sigma), including 10% FCS (Gibco), 2 mM glutamine (Thermo Fisher Scientific) and penicillin (50 Units/mL)/streptomycin (50 µg/mL) (Thermo Fisher Scientific). The slide was mounted onto an inverted microscope (Axiovert 100TV, Zeiss) equipped with a 37 °C incubator, and with an HXP 120 lamp for epifluorescence illumination, a halogen lamp for phase-contrast imaging, and a Coolsnap-HQ2 camera (Photometrics) as well as electronic shutters driven by MetaMorph software for image acquisition (Molecular Devices). Unipolar cells were chosen for analysis. Phase contrast and fluorescence images (in case of cells transfected with EGFP-tagged CapZ constructs) were acquired every 5 seconds using a ×63/1.4NA Plan apochromatic oil objective for a period of 5 min. Protrusion velocity was determined based on phase contrast-derived kymographs using MetaMorph 7.8. For every cell analyzed, kymographs from three different positions of the leading edge were obtained, independently analyzed and averaged.

#### Electron microscopy

##### Negative-stain microscopy

Immediately after SEC, the quality of the short actin filaments preparation was assessed with negative stain EM. Briefly, 4 µL of sample were deposited onto a glow-discharged-carbon-coated copper grid and incubated for 30 s. After blotting the excess sample, the grid was washed twice with KMEI buffer and stained with 0.75% (w/v) uranyl formate. The grids were subsequently imaged at 120 kV in a Tecnai Spirit microscope equipped with a LaB_6_ cathode and 4k × 4k CMOS detector F416 (TVIPS). Most importantly, we used these images to decide on the concentration used for preparing the Cryo-EM grids.

##### Cryo grid preparation

Cryo-EM samples were prepared using graphene-oxide-coated Quantifoil 1.2/1.3 300 mesh grids (GO-grids). We produce our GO-grids as follows: we took an aliquot of the 2 mg/ml stock graphene-oxide (GO) solution (Sigma-Aldrich) and sonicated it for 3 min in a bath sonicator. This step is necessary to separate aggregated GO plates. After this, we prepared dilutions between 0.05 and 0.15 mg/ml for screening. Quantifoil grids were freshly glow-discharged and 4 µL of our GO dilutions were immediately applied to the grids. After 3 min incubation, the grids were blotted from the back side using Whatmann No. 5 and washed twice with 15 µL water. The quality of the resulting GO-grids was immediately assessed in our Tecnai Spirit microscope. Typically, we looked for a compromise between grid coverage and number of GO layers, with most selected grids showing a few layers - sometimes single ones - and almost 100% grid coverage. Shortly before applying the sample, the GO-grids were mildly glow discharged for 10 s using a 5 mA current.

Vitrification was performed in Vitrobot Mark IV. Four µL of sample were placed onto a GO-grid, incubated for 1 min at 13 °C and 100% humidity, blotted for 3.5 s with a force of –3, and subsequently plunged into liquid ethane.

##### Data collection and preprocessing

Cryo-EM images were obtained using a Talos Arctica microscope equipped with a 200 kV XFEG, and automatically collected using EPU. Exposures of 3 s were recorded with a Falcon III detector operated in linear mode, as stacks of 40 frames containing a total dose of 60 e^−^/Å^2^. Due to the beam diameter of ∼1.7 µm (50 µm C2 aperture) we collected a single exposure per hole. No objective aperture was used during data collection.

We collected a total of 4,204 images in two separated sessions, covering a defocus range between –0.7 and −4.0 µm. Motion correction (Zheng et al., 2017) and CTF estimation (Rohou and Grigorieff, 2015) were performed on the fly with TranSPHIRE (Stabrin et al., 2020).

##### Cryo-EM image processing

Most image processing steps were performed using SPHIRE (Moriya et al., 2017). Filament ends were automatically detected with crYOLO (Wagner et al., 2019). In order to improve the picking, we denoised the images using the noise2noise implementation within cryolo (janni) and further filtered the images using the ctfcorr.simple. filter within the e2proc2d.py program of EMAN2 (Tang et al., 2007) (See Supplementary Figure 1). Using an ad-hoc model trained on these images we obtained a total of 570,524 initial particles. In order to separate Barbed from Pointed ends, we performed two independent runs of 3D multi-reference alignment (sp_mref_ali3d.py) against references for both ends, and kept only those that we systematically assigned to the Barbed end. The rest of the processing was performed in SPHIRE’s filament mode. This was necessary since the high density of filament images provided too many local minima for alignment, which was alleviated by the rectangular mask used by helical SPHIRE. We used the final alignment parameters obtained in the multi-reference alignment step to estimate the in-plane rotation of the filaments. We then performed 2D classification with ISAC (Yang et al., 2012) and discarded all non-protein picks, but kept all filaments, even if the class did not appear to be an end. This was necessary since, although ISAC produces validated and highly homogeneous classes, ends and full filaments could still be clearly seen mixed in some of them. With this set of clean Barbed end particles, we started a 3D refinement using a full filament as a reference, followed by focused 3D classification. In cases where the ends were out of register, we re-centered the particles, re-extracted them, and re-run 3D refinement. After a few iterations, a single class containing capping protein could be obtained. We then exported the particles to RELION (Scheres, 2012) where we performed Bayesian particle polishing (Zivanov et al., 2019), and re-imported the stack for a final round of k-means 2D clustering cleaning followed by local refinement within SPHIRE. Supplementary Figure 1 shows a schematic summary of the procedure.

##### Resolution estimation and maps postprocessing

The final map was obtained from particles with some preferential orientation. To estimate the directional FSC of the map we used the 3D-FSC package (Tan et al., 2017). Local resolution was estimated using the false-discovery rate FSC method implemented in SPOC (Beckers and Sachse, 2020). The final map was locally filtered using deepEMhancer (Sanchez-Garcia, 2020).

#### Atomic modeling

We used MODELLER (Eswar et al., 2008) to build a homology model of the complex between CP and the barbed end of human β-actin. As template, we used the capped end of the Arp1 filaments within the dynactin complex (PDBID:5ADX, (Urnavicius et al., 2015)). We run this model through the Rosetta alanine scan server (Kortemme et al., 2004) to identify side chains whose deletion would provide a similar decrease in binding affinity as the removal of the β tentacle.

To build an atomic model of the complex, with fitted the initial homology model into the density using a combination of ISOLDE (Croll, 2018) and Coot (Emsley et al., 2010). During Isolde’s flexible fitting, we always included distance restraints aimed at keeping the structure of CP in place. This was necessary due to the moderate resolution of the map in this area. A final round of refinement in Phenix was used to remove some geometric violation and estimate the atomic B-factors.

#### Quantification and statistical data analysis

All analyzed data were plotted and fitted in Origin9.0G. All fluorescence microscopy experiments were analyzed in ImageJ manually via kymograph analysis and plotting of intensity profiles unless described otherwise.

##### Determination of the association and dissociation rate constants of capping protein for actin filament ends

Images were analyzed by manual filament tracking using the *segmented line tool* from ImageJ and further analyzed by the *kymograph plugin*. The slopes were measured to determine the polymerization rate of individual actin filaments. The pixel size/length was converted into microns/s. One actin monomer contributes to 2.7 nm of the actin filament length. For each experimental condition, the filament polymerization velocity was measured from ≥50 filaments per condition, and reported as mean value with error bars representing SD.

The time course [sec] of the fraction of capped filament ends was fitted with a mono-exponential function to yield the observed pseudo-first order reaction rate (k_obs_) as a function of capping protein concentration. The association rate constants (k_on_) were determined from linear regression fits of the k_obs_ mean values (error = SD) as a function of the total capping protein concentration [nM].

The time course [sec] of the fraction of re-growing filament ends was fitted with a single exponential function to yield the dissociation rate constant (k_off_).

The equilibrium dissociation constants (K_D_) of capping protein for actin filament barbed ends were calculated from the measured dissociation rate constants (k_off_) and association rate constants (k_on_) using the following equation:

1. *KD*=*koff / kon*

Errors for the equilibrium dissociation constants were calculated using error propagation.

##### Quantification of fluorescence intensities in actin dendritic networks

The mean fluorescence intensities of all network components (actin, CP and Arp2/3) from multicolor fluorescence images were measured with ImageJ (ROI Manager->Multi Measure function) from line regions of interest (ROIs) matching the network area. The background fluorescence intensities for all components was determined from identical lines drawn adjacent to each network. The background signal was subtracted from the network signal. The integrated mean fluorescence intensities (representing polymerization, capping and nucleation rates) for each component were plotted as a function of distance from the bead surface [μm]. The rates for actin polymerization, capping and nucleation were calculated from integrated fluorescence intensities divided by reaction time.

##### Quantification of the NPF-WH2-occupancy on actin filament ends

The fluorescence signal intensity of the Alexa488-labeled NPF was measured over the entire bead surface before and after addition of quencher-labeled Atto540Q actin monomers. The intensity data was plotted over the bead distance/diameter and represents the amount of WH_2_ domains occupied by actin filament ends. For each experiment, ≥ 25 dendritic actin networks from ≥ 3 slides were quantified.

## References

Achard V, Martiel J-L, Michelot A, Guérin C, Reymann A-C, Blanchoin L, Boujemaa-Paterski R. 2010. A “primer”-based mechanism underlies branched actin filament network formation and motility. Curr Biol 20:423–428. doi:10.1016/j.cub.2009.12.056

Akin O, Mullins RD. 2008. Capping protein increases the rate of actin-based motility by promoting filament nucleation by the Arp2/3 complex. Cell 133:841–851. doi:10.1016/j.cell.2008.04.011

Balzer CJ, Wagner AR, Helgeson LA, Nolen BJ. 2018. Dip1 Co-opts Features of Branching Nucleation to Create Linear Actin Filaments that Activate WASP-Bound Arp2/3 Complex. Curr Biol 28:3886–3891.e4. doi:10.1016/j.cub.2018.10.045

Bieling P, Hansen SD, Akin O, Li T-D, Hayden CC, Fletcher DA, Mullins RD. 2018. WH2 and proline-rich domains of WASP-family proteins collaborate to accelerate actin filament elongation. EMBO J 37:102–121. doi:10.15252/embj.201797039

Bieling P, Li T-D, Weichsel J, McGorty R, Jreij P, Huang B, Fletcher DA, Mullins RD. 2016. Force Feedback Controls Motor Activity and Mechanical Properties of Self-Assembling Branched Actin Networks. Cell 164:115–127. doi:10.1016/j.cell.2015.11.057

Blanchoin L, Boujemaa-Paterski R, Sykes C, Plastino J. 2014. Actin dynamics, architecture, and mechanics in cell motility. Physiol Rev 94:235–263. doi:10.1152/physrev.00018.2013

Bombardier JP, Eskin JA, Jaiswal R, Corrêa IR, Xu M-Q, Goode BL, Gelles J. 2015. Single-molecule visualization of a formin-capping protein “decision complex” at the actin filament barbed end. Nat Commun 6:8707. doi:10.1038/ncomms9707

Burkel BM, von Dassow G, Bement WM. 2007. Versatile fluorescent probes for actin filaments based on the actin-binding domain of utrophin. Cell Motil Cytoskeleton 64:822–832. doi:10.1002/cm.20226

Cameron LA, Svitkina TM, Vignjevic D, Theriot JA, Borisy GG. 2001. Dendritic organization of actin comet tails. Curr Biol 11:130–135. doi:10.1016/s0960-9822(01)00022-7

Chereau D, Kerff F, Graceffa P, Grabarek Z, Langsetmo K, Dominguez R. 2005. Actin-bound structures of Wiskott-Aldrich syndrome protein (WASP)-homology domain 2 and the implications for filament assembly. Proc Natl Acad Sci U S A 102:16644–16649. doi:10.1073/pnas.0507021102

Co C, Wong DT, Gierke S, Chang V, Taunton J. 2007. Mechanism of actin network attachment to moving membranes: barbed end capture by N-WASP WH2 domains. Cell 128:901–913. doi:10.1016/j.cell.2006.12.049

Cooper JA, Pollard TD. 1985. Effect of capping protein on the kinetics of actin polymerization. Biochemistry 24:793–799. doi:10.1021/bi00324a039

Cooper JA, Sept D. 2008. New insights into mechanism and regulation of actin capping protein. Int Rev Cell Mol Biol 267:183–206. doi:10.1016/S1937-6448(08)00604-7

Edwards M, Zwolak A, Schafer DA, Sept D, Dominguez R, Cooper JA. 2014. Capping protein regulators fine-tune actin assembly dynamics. Nat Rev Mol Cell Biol 15:677–689. doi:10.1038/nrm3869

Efimova N, Svitkina TM. 2018. Branched actin networks push against each other at adherens junctions to maintain cell-cell adhesion. J Cell Biol 217:1827–1845. doi:10.1083/jcb.201708103

Funk J, Merino F, Venkova L, Heydenreich L, Kierfeld J, Vargas P, Raunser S, Piel M, Bieling P. 2019. Profilin and formin constitute a pacemaker system for robust actin filament growth. Elife 8. doi:10.7554/eLife.50963

Hakala M, Wioland H, Tolonen M, Kotila T, Jegou A, Romet-Lemonne G, Lappalainen P. 2021. Twinfilin uncaps filament barbed ends to promote turnover of lamellipodial actin networks. Nat Cell Biol 23:147–159. doi:10.1038/s41556-020-00629-y

Hart MC, Korshunova YO, Cooper JA. 1997. Vertebrates have conserved capping protein α isoforms with specific expression patterns. Cell Motility and the Cytoskeleton 38:120–132. doi:10.1002/(SICI)1097-0169(1997)38:2<120::AID-CM2>3.0.CO;2-B

Helgeson LA, Nolen BJ. 2013. Mechanism of synergistic activation of Arp2/3 complex by cortactin and N-WASP. Elife 2:e00884. doi:10.7554/eLife.00884

Hug C, Jay PY, Reddy I, McNally JG, Bridgman PC, Elson EL, Cooper JA. 1995. Capping protein levels influence actin assembly and cell motility in dictyostelium. Cell 81:591–600. doi:10.1016/0092-8674(95)90080-2

Hurst S, Howes EA, Coadwell J, Jones R. 1998. Expression of a testis-specific putative actin-capping protein associated with the developing acrosome during rat spermiogenesis. Mol Reprod Dev 49:81–91. doi:10.1002/(SICI)1098-2795(199801)49:1<81::AID-MRD9>3.0.CO;2-K

Isenberg G, Aebi U, Pollard TD. 1980. An actin-binding protein from Acanthamoeba regulates actin filament polymerization and interactions. Nature 288:455–459. doi:10.1038/288455a0

Iwasa JH, Mullins RD. 2007. Spatial and temporal relationships between actin-filament nucleation, capping, and disassembly. Curr Biol 17:395–406. doi:10.1016/j.cub.2007.02.012

Jaumouillé V, Waterman CM. 2020. Physical Constraints and Forces Involved in Phagocytosis. Front Immunol 11:1097. doi:10.3389/fimmu.2020.01097

Kaiser DA, Vinson VK, Murphy DB, Pollard TD. 1999. Profilin is predominantly associated with monomeric actin in Acanthamoeba. J Cell Sci 112 (Pt 21):3779–3790.

Kast DJ, Dominguez R. 2017. The Cytoskeleton-Autophagy Connection. Curr Biol 27:R318–R326. doi:10.1016/j.cub.2017.02.061

Kim K, Yamashita A, Wear MA, Maéda Y, Cooper JA. 2004. Capping protein binding to actin in yeast: biochemical mechanism and physiological relevance. J Cell Biol 164:567–580. doi:10.1083/jcb.200308061

Kim T, Cooper JA, Sept D. 2010. The interaction of capping protein with the barbed end of the actin filament. J Mol Biol 404:794–802. doi:10.1016/j.jmb.2010.10.017

Lai FPL, Szczodrak M, Block J, Faix J, Breitsprecher D, Mannherz HG, Stradal TEB, Dunn GA, Small JV, Rottner K. 2008. Arp2/3 complex interactions and actin network turnover in lamellipodia. EMBO J 27:982–992. doi:10.1038/emboj.2008.34

Loisel TP, Boujemaa R, Pantaloni D, Carlier MF. 1999. Reconstitution of actin-based motility of Listeria and Shigella using pure proteins. Nature 401:613–616. doi:10.1038/44183

Marchand JB, Kaiser DA, Pollard TD, Higgs HN. 2001. Interaction of WASP/Scar proteins with actin and vertebrate Arp2/3 complex. Nat Cell Biol 3:76–82. doi:10.1038/35050590

Mejillano MR, Kojima S, Applewhite DA, Gertler FB, Svitkina TM, Borisy GG. 2004. Lamellipodial versus filopodial mode of the actin nanomachinery: pivotal role of the filament barbed end. Cell 118:363–373. doi:10.1016/j.cell.2004.07.019

Merino F, Pospich S, Funk J, Wagner T, Küllmer F, Arndt H-D, Bieling P, Raunser S. 2018. Structural transitions of F-actin upon ATP hydrolysis at near-atomic resolution revealed by cryo-EM. Nat Struct Mol Biol 25:528–537. doi:10.1038/s41594-018-0074-0

Miyoshi T, Tsuji T, Higashida C, Hertzog M, Fujita A, Narumiya S, Scita G, Watanabe N. 2006. Actin turnover-dependent fast dissociation of capping protein in the dendritic nucleation actin network: evidence of frequent filament severing. J Cell Biol 175:947–955. doi:10.1083/jcb.200604176

Mueller J, Szep G, Nemethova M, de Vries I, Lieber AD, Winkler C, Kruse K, Small JV, Schmeiser C, Keren K, Hauschild R, Sixt M. 2017. Load Adaptation of Lamellipodial Actin Networks. Cell 171:188–200.e16. doi:10.1016/j.cell.2017.07.051

Mullins RD, Bieling P, Fletcher DA. 2018. From solution to surface to filament: actin flux into branched networks. Biophys Rev 10:1537–1551. doi:10.1007/s12551-018-0469-5

Mwangangi DM, Manser E, Robinson RC. 2021. The structure of the actin filament uncapping complex mediated by twinfilin. Sci Adv 7. doi:10.1126/sciadv.abd5271

Narita A, Takeda S, Yamashita A, Maéda Y. 2006. Structural basis of actin filament capping at the barbed-end: a cryo-electron microscopy study. EMBO J 25:5626–5633. doi:10.1038/sj.emboj.7601395

Oda T, Iwasa M, Aihara T, Maéda Y, Narita A. 2009. The nature of the globular- to fibrous-actin transition. Nature 457:441–445. doi:10.1038/nature07685

Papalazarou V, Machesky LM. 2020. The cell pushes back: The Arp2/3 complex is a key orchestrator of cellular responses to environmental forces. Curr Opin Cell Biol 68:37–44. doi:10.1016/j.ceb.2020.08.012

Pospich S, Merino F, Raunser S. 2020. Structural Effects and Functional Implications of Phalloidin and Jasplakinolide Binding to Actin Filaments. Structure 28:437–449.e5. doi:10.1016/j.str.2020.01.014

Prostak SM, Robinson KA, Titus MA, Fritz-Laylin LK. 2020. The actin networks of chytrid fungi reveal evolutionary loss of cytoskeletal complexity in the fungal kingdom (preprint). Cell Biology. doi:10.1101/2020.06.12.142943

Rivero F, Cvrcková F. 2007. Origins and evolution of the actin cytoskeleton. Adv Exp Med Biol 607:97–110. doi:10.1007/978-0-387-74021-8_8

Rotty JD, Wu C, Bear JE. 2013. New insights into the regulation and cellular functions of the ARP2/3 complex. Nat Rev Mol Cell Biol 14:7–12. doi:10.1038/nrm3492

Sanchez-Garcia R, Gomez-Blanco J, Cuervo A, Carazo J, Sorzano C, Vargas J. 2020. DeepEMhancer: a deep learning solution for cryo-EM volume post-processing (preprint). Bioinformatics. doi:10.1101/2020.06.12.148296

Schafer DA, Jennings PB, Cooper JA. 1996. Dynamics of capping protein and actin assembly in vitro: uncapping barbed ends by polyphosphoinositides. J Cell Biol 135:169–179. doi:10.1083/jcb.135.1.169

Schafer DA, Korshunova YO, Schroer TA, Cooper JA. 1994. Differential localization and sequence analysis of capping protein beta-subunit isoforms of vertebrates. J Cell Biol 127:453–465. doi:10.1083/jcb.127.2.453

Senju Y, Lappalainen P. 2019. Regulation of actin dynamics by PI(4,5)P2 in cell migration and endocytosis. Curr Opin Cell Biol 56:7–13. doi:10.1016/j.ceb.2018.08.003

Shaaban M, Chowdhury S, Nolen BJ. 2020. Cryo-EM reveals the transition of Arp2/3 complex from inactive to nucleation-competent state. Nat Struct Mol Biol 27:1009–1016. doi:10.1038/s41594-020-0481-x

Shekhar S, Kerleau M, Kühn S, Pernier J, Romet-Lemonne G, Jégou A, Carlier M-F. 2015. Formin and capping protein together embrace the actin filament in a ménage à trois. Nat Commun 6:8730. doi:10.1038/ncomms9730

Urnavicius L, Zhang K, Diamant AG, Motz C, Schlager MA, Yu M, Patel NA, Robinson CV, Carter AP. 2015. The structure of the dynactin complex and its interaction with dynein. Science 347:1441–1446. doi:10.1126/science.aaa4080

Vinzenz M, Nemethova M, Schur F, Mueller J, Narita A, Urban E, Winkler C, Schmeiser C, Koestler SA, Rottner K, Resch GP, Maeda Y, Small JV. 2012. Actin branching in the initiation and maintenance of lamellipodia. J Cell Sci 125:2775–2785. doi:10.1242/jcs.107623

Wang R, Carlsson AE. 2015. How capping protein enhances actin filament growth and nucleation on biomimetic beads. Phys Biol 12:066008. doi:10.1088/1478-3975/12/6/066008

Wear MA, Yamashita A, Kim K, Maéda Y, Cooper JA. 2003. How capping protein binds the barbed end of the actin filament. Curr Biol 13:1531–1537. doi:10.1016/s0960-9822(03)00559-1

Wu C, Asokan SB, Berginski ME, Haynes EM, Sharpless NE, Griffith JD, Gomez SM, Bear JE. 2012. Arp2/3 is critical for lamellipodia and response to extracellular matrix cues but is dispensable for chemotaxis. Cell 148:973–987. doi:10.1016/j.cell.2011.12.034

Ydenberg CA, Smith BA, Breitsprecher D, Gelles J, Goode BL. 2011. Cease-fire at the leading edge: new perspectives on actin filament branching, debranching, and cross-linking. Cytoskeleton (Hoboken) 68:596–602. doi:10.1002/cm.20543

## Supplemental References

Aitken, C.E., Marshall, R.A., and Puglisi, J.D. (2008). An oxygen scavenging system for improvement of dye stability in single-molecule fluorescence experiments. Biophys J 94, 1826–1835.

Akin, O., and Mullins, R.D. (2008). Capping protein increases the rate of actin-based motility by promoting filament nucleation by the Arp2/3 complex. Cell 133, 841–851.

Antos, J.M., Ingram, J., Fang, T., Pishesha, N., Truttmann, M.C., and Ploegh, H.L. (2017). Site-Specific Protein Labeling via Sortase-Mediated Transpeptidation. Curr Protoc Protein Sci 89, 15 13 11–15 13 19.

Beckers, M., and Sachse, C. (2020). Permutation testing of Fourier shell correlation for resolution estimation of cryo-EM maps. J Struct Biol 212, 107579.

Bieling, P., Hansen, S.D., Akin, O., Li, T.D., Hayden, C.C., Fletcher, D.A., and Mullins, R.D. (2018). WH2 and proline-rich domains of WASP-family proteins collaborate to accelerate actin filament elongation. EMBO J 37, 102–121.

Bieling, P., Li, T.D., Weichsel, J., McGorty, R., Jreij, P., Huang, B., Fletcher, D.A., and Mullins, R.D. (2016). Force Feedback Controls Motor Activity and Mechanical Properties of Self-Assembling Branched Actin Networks. Cell 164, 115–127.

Bieling, P., Telley, I.A., Hentrich, C., Piehler, J., and Surrey, T. (2010). Fluorescence microscopy assays on chemically functionalized surfaces for quantitative imaging of microtubule, motor, and +TIP dynamics. Methods Cell Biol 95, 555–580.

Brinkman, E.K., Chen, T., Amendola, M., and van Steensel, B. (2014). Easy quantitative assessment of genome editing by sequence trace decomposition. Nucleic Acids Res 42, e168.

Croll, T.I. (2018). ISOLDE: a physically realistic environment for model building into low-resolution electron-density maps. Acta Crystallogr D Struct Biol 74, 519–530.

Doolittle, L.K., Rosen, M.K., and Padrick, S.B. (2013). Purification of native Arp2/3 complex from bovine thymus. Methods Mol Biol 1046, 231–250.

Emsley, P., Lohkamp, B., Scott, W.G., and Cowtan, K. (2010). Features and development of Coot. Acta Crystallogr D Biol Crystallogr 66, 486–501.

Eswar, N., Eramian, D., Webb, B., Shen, M.Y., and Sali, A. (2008). Protein structure modeling with MODELLER. Methods Mol Biol 426, 145–159.

Funk, J., Merino, F., Venkova, L., Heydenreich, L., Kierfeld, J., Vargas, P., Raunser, S., Piel, M., and Bieling, P. (2019). Profilin and formin constitute a pacemaker system for robust actin filament growth. Elife 8.

Gibson, D.G. (2011). Enzymatic assembly of overlapping DNA fragments. Methods Enzymol 498, 349–361.

Hansen, S.D., and Mullins, R.D. (2010). VASP is a processive actin polymerase that requires monomeric actin for barbed end association. J Cell Biol 191, 571–584.

Hansen, S.D., Zuchero, J.B., and Mullins, R.D. (2013). Cytoplasmic actin: purification and single molecule assembly assays. Methods Mol Biol 1046, 145–170.

Keppler, A., Gendreizig, S., Gronemeyer, T., Pick, H., Vogel, H., and Johnsson, K. (2003). A general method for the covalent labeling of fusion proteins with small molecules in vivo. Nat Biotechnol 21, 86–89.

Kortemme, T., Kim, D.E., and Baker, D. (2004). Computational alanine scanning of protein-protein interfaces. Sci STKE 2004, pl2.

Kuhn, J.R., and Pollard, T.D. (2005). Real-time measurements of actin filament polymerization by total internal reflection fluorescence microscopy. Biophys J 88, 1387–1402.

Moriya, T., Saur, M., Stabrin, M., Merino, F., Voicu, H., Huang, Z., Penczek, P.A., Raunser, S., and Gatsogiannis, C. (2017). High-resolution Single Particle Analysis from Electron Cryo-microscopy Images Using SPHIRE. J Vis Exp.

Pollard, T.D. (1982). Myosin purification and characterization. Methods Cell Biol 24, 333–371.

Rasnik, I., McKinney, S.A., and Ha, T. (2006). Nonblinking and long-lasting single-molecule fluorescence imaging. Nat Methods 3, 891–893.

Rohou, A., and Grigorieff, N. (2015). CTFFIND4: Fast and accurate defocus estimation from electron micrographs. J Struct Biol 192, 216–221.

Sanchez-Garcia, R., J. Gomez-Blanco, A. Cuervo, J. M. Carazo, C. O. S. Sorzano, and J. Vargas (2020). DeepEMhancer: A Deep Learning Solution for Cryo-EM Volume Post-Processing. BioRxiv.

Scheres, S.H. (2012). RELION: implementation of a Bayesian approach to cryo-EM structure determination. J Struct Biol 180, 519–530.

Stabrin, M., Schoenfeld, F., Wagner, T., Pospich, S., Gatsogiannis, C., and Raunser, S. (2020). TranSPHIRE: automated and feedback-optimized on-the-fly processing for cryo-EM. Nat Commun 11, 5716.

Tan, Y.Z., Baldwin, P.R., Davis, J.H., Williamson, J.R., Potter, C.S., Carragher, B., and Lyumkis, D. (2017). Addressing preferred specimen orientation in single-particle cryo-EM through tilting. Nat Methods 14, 793–796.

Tang, G., Peng, L., Baldwin, P.R., Mann, D.S., Jiang, W., Rees, I., and Ludtke, S.J. (2007). EMAN2: an extensible image processing suite for electron microscopy. J Struct Biol 157, 38–46.

Urnavicius, L., Zhang, K., Diamant, A.G., Motz, C., Schlager, M.A., Yu, M., Patel, N.A., Robinson, C.V., and Carter, A.P. (2015). The structure of the dynactin complex and its interaction with dynein. Science 347, 1441–1446.

Wagner, T., Merino, F., Stabrin, M., Moriya, T., Antoni, C., Apelbaum, A., Hagel, P., Sitsel, O., Raisch, T., Prumbaum, D., et al. (2019). SPHIRE-crYOLO is a fast and accurate fully automated particle picker for cryo-EM. Commun Biol 2, 218.

Yang, Z., Fang, J., Chittuluru, J., Asturias, F.J., and Penczek, P.A. (2012). Iterative stable alignment and clustering of 2D transmission electron microscope images. Structure 20, 237–247.

Zheng, S.Q., Palovcak, E., Armache, J.P., Verba, K.A., Cheng, Y., and Agard, D.A. (2017). MotionCor2: anisotropic correction of beam-induced motion for improved cryo-electron microscopy. Nat Methods 14, 331–332.

Zivanov, J., Nakane, T., and Scheres, S.H.W. (2019). A Bayesian approach to beam-induced motion correction in cryo-EM single-particle analysis. IUCrJ 6, 5–17.

